# Brain Landscape In Situ Crosslinking Mass Spectrometry (BLIS-XL-MS) Enables Global Analysis of Protein Structural Remodeling in the Brain

**DOI:** 10.64898/2026.09.16.752161

**Authors:** Yi-Zhi Wang, Jian Xu, Anis Contractor, Jeffrey N. Savas

## Abstract

Pathological protein conformational remodeling is a key molecular feature of brain disorders. However, conventional approaches to understand the changes in potein conformation are targeted and difficult to scale proteome-wide analysis. Emerging mass spectrometry-based structural proteomic methods broaden coverage but often involve complex workflows and trade-offs between labeling efficiency and preservation of the *in vivo* molecular state. Thus, quantitative analysis of proteome-wide structural remodeling remains limited. Here we present Brain Landscape In Situ Crosslinking Mass Spectrometry (BLIS-XL-MS), an experimental and computational framework for analyzing protein and protein-complex remodeling in brain tissue. BLIS-XL-MS stabilizes the molecular state before secondary crosslinking, improves reagent accessibility through sectioning and permeabilization, and integrates quantitative crosslink analysis with structural and protein-interaction-network interpretation. Applied to brains from GluA1^A636T^ knock-in mice, modeling a neurodevelopmental disorder, BLIS-XL-MS revealed coordinated remodeling of protein systems involved in AMPA receptor-trafficking, mitochondrial function and cell-death pathways. We demonstrate that BLIS-XL-MS provides a scalable framework for mapping disease-associated structural remodeling across the brain proteome.

## Main

Protein conformation is a fundamental determinant of proper protein function, molecular interactions, and complex assembly^1^. In many brain disorders, molecular pathology extends beyond changes in protein abundance and post-translational modifications (PTMs) to include alterations in protein conformation, folding, assembly into complexes, and interaction states^2–4^. Defining these conformational changes can therefore uncover disease mechanisms and therapeutic opportunities that are not revealed by abundance-based proteomics. However, most conventional approaches for studying protein conformation, including cryo-electron microscopy (Cryo-EM), nuclear magnetic resonance spectroscopy, and Förster resonance energy transfer, are targeted to individual proteins or complexes rather than applied proteome-wide. These approaches are difficult to scale for unbiased, proteome-wide analysis of protein structural remodeling in complex tissues.

Mass spectrometry (MS)-based structural proteomics, including limited proteolysis, covalent footprinting and crosslinking MS (XL-MS), offer proteome-wide readouts^5–7^. However, most implementations have analyzed on tissue homogenates, cultured cells, or isolated subcellular fractions organelles, or affinity purified protein complexes. Sample preparation steps such as homogenization and fractionation disrupt tissue architecture and compartmentalization while altering molecular concentrations, macromolecular crowding, membrane organization, and local interaction environments^8–10^. Indeed, tissue deuterium-exchange MS revealed that the same proteins exhibit distinct conformational dynamics in *ex vivo* brain sections compared with soluable extracts^11^.

Recent advances in XL-MS are starting to over come these limitations and direct tissue XL-MS has been applied to freshly dissected heart and skeletal muscle^12–15^. These approaches preserve tissue context more effectively than post-lysis crosslinking. However, in fresh brain tissue, conditions that improve reagent penetration can also disrupt native molecular states, creating a trade-off between tissue accessibility and preservation of protein structure. Brain dissection rapidly perturbs neuronal metabolism, ionic homeostasis, signaling and synaptic architecture, raising the concern that prolonged *ex vivo* crosslinking captures progressively remodeled protein conformational and interaction states rather than the original *in vivo* molecular state. Limited crosslinker diffusion through dense, lipid-rich brain tissue can prolong this window, whereas longer incubations, high reagent concentrations and organic cosolvents, such as DMSO, further perturb protein conformations and interactions. Consequently, slow in-tissue crosslinking can capture a time-integrated mixture of native and post-dissection states, underscoring the need to arrest the brain molecular state before secondary crosslinking. Thus, conditions that have previously been demonstrated to provide adequate crosslinking coverage in fresh heart or skeletal muscle cannot be assumed to preserve the molecular state of fresh brain tissue, creating a critical need for devising brain-specific in-tissue XL-MS strategies that rapidly arrest molecular dynamics, minimize diffusion distance and enable efficient crosslinker access.

Moreover, comparative analysis across XL-MS datasets presents an additional challenge. XL-MS datasets contain looplinks where the crosslinker connects residues in the same peptide, intralinks where the crosslinker will connect two different peptides in the same protein and interlinks where crosslinking occurs between peptides in two separate proteins. Quantitative XL-MS commonly evaluates structural differences through changes in the abundance or detection of individual crosslink or residue pairs. However, data-dependent acquisition (DDA), which remains the predominant acquisition strategy in XL-MS, incompletely samples the available precursor population, with intensity-dependent selection and stochastic variation between runs. Consequently, failure to identify a crosslink does not establish the absence of the corresponding molecular proximity or interaction. Comparisons that rely exclusively on condition-specific detections or a small number of sparsely supported crosslinks can therefore conflate sampling variability with biological remodeling. Therefore, we need to profile and compare the distribution of crosslink evidence within each protein, rather than relying solely on isolated crosslink changes. Similarly, at the protein-complex level, we need to evaluate coordinated changes in interaction organization within consistently defined modules, rather than interpreting individual crosslink gains or losses in isolation.

Here we introduce Brain Landscape In Situ Crosslinking Mass Spectrometry (BLIS-XL-MS), comprising a brain tissue-compatible experimental workflow and the BLIS 1.0 computational package (**Fig. 1**). BLIS-XL-MS combines paraformaldehyde (PFA) stabilization, vibratrome sectioning, membrane permeabilization and MS-cleavable crosslinker disuccinimidyl sulfoxide (DSSO). This design fixes the brain tissue before protein crosslinking, shortens diffusion distance and improves intracellular access. BLIS 1.0 integrates quality-weighted crosslink evidence into protein-level structural profiles and condition-blind protein-protein interaction (PPI) modules to quantify condition-associated remodeling from individual proteins to protein complexes and interaction networks. We applied the framework to adult heterozygous GluA1^A636T^ knock-in mice carrying a pathogenic GluA1 gating variant associated with neurodevelopmental disorder (NDD). BLIS-XL-MS identified remodeling of key regulators of AMPAR trafficking and mitochondrial Complex V in the brain of A636T mice. These signatures converged with abnormalities independently identified previously by electrophysiological, quantitative proteomic, imaging and histopathological measurements in these mice^16,17^. BLIS-XL-MS therefore provides a brain tissue-compatible framework for comparative structural systems biology, generating structural and interaction-level hypotheses that complement conventional abundance-based proteomics and orthogonal functional and pathological measurements.

**Fig. 1.**
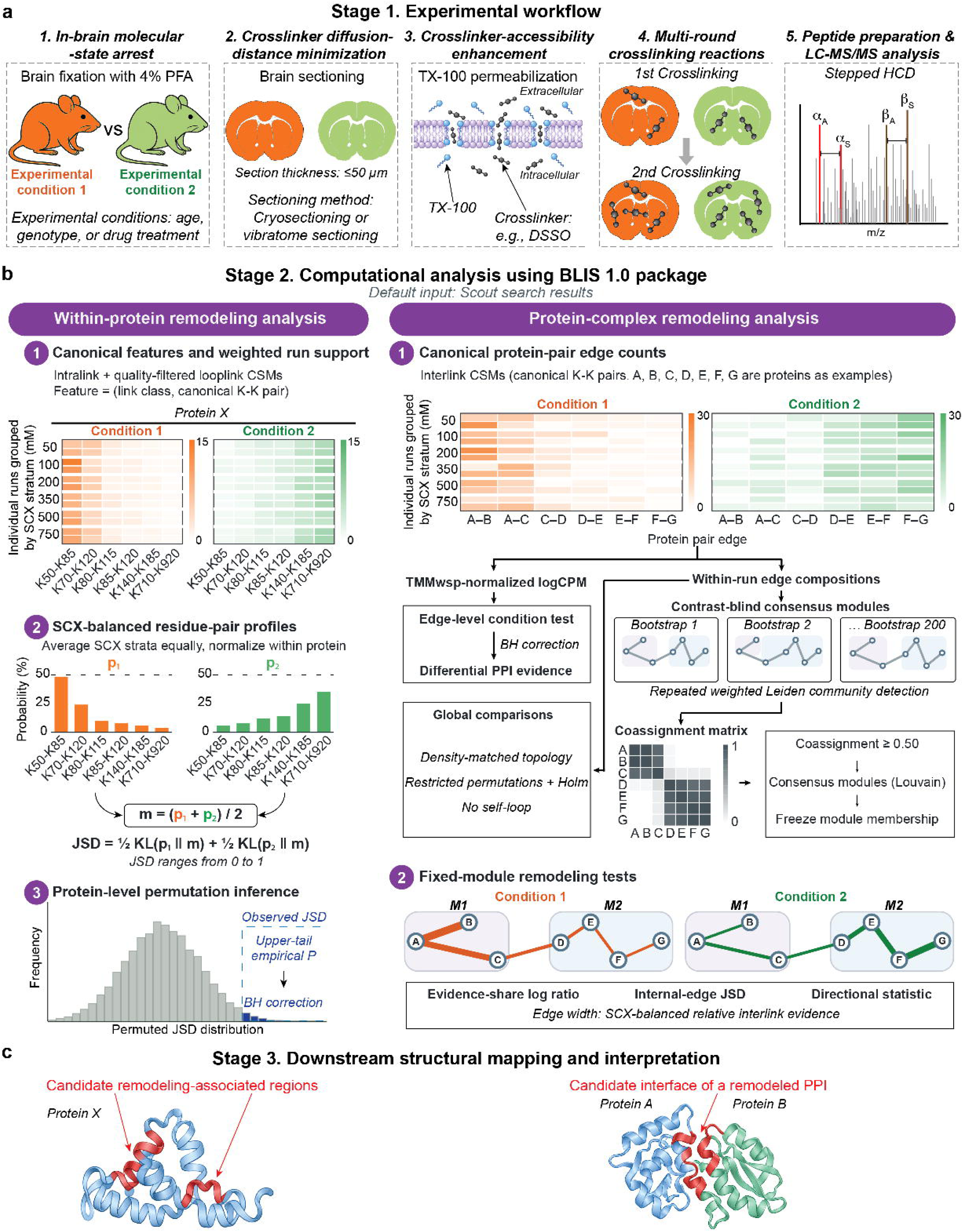
BLIS-XL-MS experimental and computational framework for proteome-scale analysis of within-protein conformational and protein complex remodeling in brain tissue. **a,** Stage 1, experimental workflow of BLIS-XL-MS. Brains from animals representing two experimental conditions, such as different ages, drug treatments or genotypes, are fixed with 4% PFA to arrest the tissue molecular state, followed by sectioning, permeabilization, secondary crosslinking with DSSO and quenching. PFA-derived crosslinks are subsequently reversed during sample processing, and the tissue is homogenized. Proteins are reduced and alkylated, purified by single-pot solid-phase-enhanced sample preparation (SP3), sequentially digested with LysC and trypsin, desalted, and fractionated by orthogonal SCX and high-pH reversed-phase separation before stepped-HCD LC-MS/MS analysis. **b,** Stage 2, BLIS 1.0 computational analysis. Crosslinked peptides are identified with Scout and filtered at the CSM, residue-pair and PPI levels. Within-protein remodeling integrates intralinks with quality-filtered looplinks, maps crosslinked lysines to canonical coordinates, and summarizes evidence as condition-specific weighted residue-pair profiles. Profile differences are quantified by weighted Jensen-Shannon divergence (JSD), followed by stratified permutation testing and BH-FDR correction to rank remodeled protein candidates. Protein-complex remodeling maps interlinks to canonical protein pairs to construct normalized edge-by-run interaction matrices and evaluates quantitative edge changes, condition-restricted occupancy, interaction composition and network topology. Condition-blind consensus modules are derived from the combined network, and restricted permutation testing with analysis-specific multiple-testing correction identifies remodeled PPI edges, candidate interfaces, network modules and topological changes. Please also see **Extended Data Fig.1a-c and Methods**. **c,** Stage 3, structural assessment of prioritized remodeling candidates. Selected proteins and complexes are evaluated using AlphaFold 3 to generate plausible structural models and assess model confidence. Where appropriate, AlphaLink2 is additionally used to incorporate experimentally observed crosslink restraints into structure prediction and to evaluate XL- supported candidate architectures. These predicted structures provide a framework for localizing BLIS remodeling signals to candidate domains, interfaces and structural regions.

## Results

### BLIS-XL-MS enables in-tissue comparative structural proteomics in the brain

Recently, a formaldehyde-based decoupled fixation and secondary-crosslinking strategy was established in cultured cells^18^. Here, we adapted this concept and further improved to address diffusion and intracellular-access constraints unique to intact brain tissue (**Fig. 1a**). Freshly prepared 4% PFA stabilized tissue architecture and molecular proximities before vibratome sectioning, Triton X-100 permeabilization and DSSO crosslinking. Unlike direct crosslinking of fresh tissue, this workflow confines the principal state-capture interval to dissection and fixation, reducing continued *ex vivo* remodeling during the longer DSSO reaction. Brain sectioning (50 μM) shortens the distance that extracellular DSSO must diffuse to reach intracellular proteins. Triton X-100 permeabilization further improves the access of crosslinkers with limited membrane permeability, such as DSSO, to intracellular structures, promoting more uniform proteome-wide crosslinking.

BLIS 1.0 first requires XL-MS raw files to be searched using Scout and subsequently takes Scout report files as the default input for downstream analyses (**Fig. 1b & Extended Data Fig.1**)^19^. BLIS 1.0 integrates quality-filtered, canonically mapped intralink and looplink evidence into protein-specific residue-pair profiles, incorporating confidence weighting and balancing across strong-cation-exchange (SCX) strata (**Extended Data Fig.1**). These profiles summarize the distribution of retained crosslink evidence within each protein, rather than relying solely on isolated crosslink changes. Differences between conditions are quantified using Jensen-Shannon divergence (JSD), with statistical support assessed by SCX-restricted whole-run label permutations and multiple-testing correction ^20,21^.

At the protein-complex level, interlink evidence is aggregated into protein-pair edges and analyzed both individually and collectively. Bootstrap-based, contrast-blind consensus modules provide a shared organizational framework for assessing coordinated remodeling: modules are defined without using observed differential edge effects or significance values, and their membership is held fixed during condition comparisons. This approach evaluates changes in interaction organization within consistently defined modules, rather than interpreting individual crosslink gains or losses in isolation.

Futhermore, AlphaFold 3 and, where appropriate, XL-restrained AlphaLink2 modeling provide structural hypotheses for prioritized candidates (**Fig. 1c**)^22,23^. These models are used to identify candidate remodeling-associated protein regions and PPI interfaces, and to prioritize residues and interaction surfaces for targeted follow-up using conventional structural approaches, such as cryo-EM, as well as biological validation and structure-guided development of molecular probes or PPI modulators.

### Example analysis of adult *Gria1* A636T heterozygous and littermate control mouse brains

A636T is a gain-of-function mutation within the highly conserved SYTANLAAF gating motif of the GluA1 subunit that enhances channel opening and markedly impairs AMPA receptor (AMPAR) desensitization^24^. GluA1^A636T^ mice exhibit deficits in hippocampal-dependent learning and memory, together with other autism spectrum disorder (ASD) and intellectual disability (ID)- relevant behavioral phenotypes^16,17,25^. However, the molecular mechanisms by which this mutation drives these brain-level abnormalities remain poorly understood.

To demonstrate the capabilities of BLIS-XL-MS, we analyzed one 3-month-old male GluA1^A636T^ mouse and one WT littermate (**Fig. 2a**). Brains were post-fixed overnight in 4% PFA at 4°C and sectioned on a vibratome at 50-µm thickness. Ten sections spanning the major hippocampal regions were analyzed per brain (**Extended Data Fig.2a**). Scout searches identified 107,770 looplinks, 7,356 intralinks and 1,516 interlinks in GluA1^A636T^ tissue, compared with 119,163 looplinks, 4,450 intralinks and 1,196 interlinks in WT control tissue, filtering at a < 1%

**Fig. 2.**
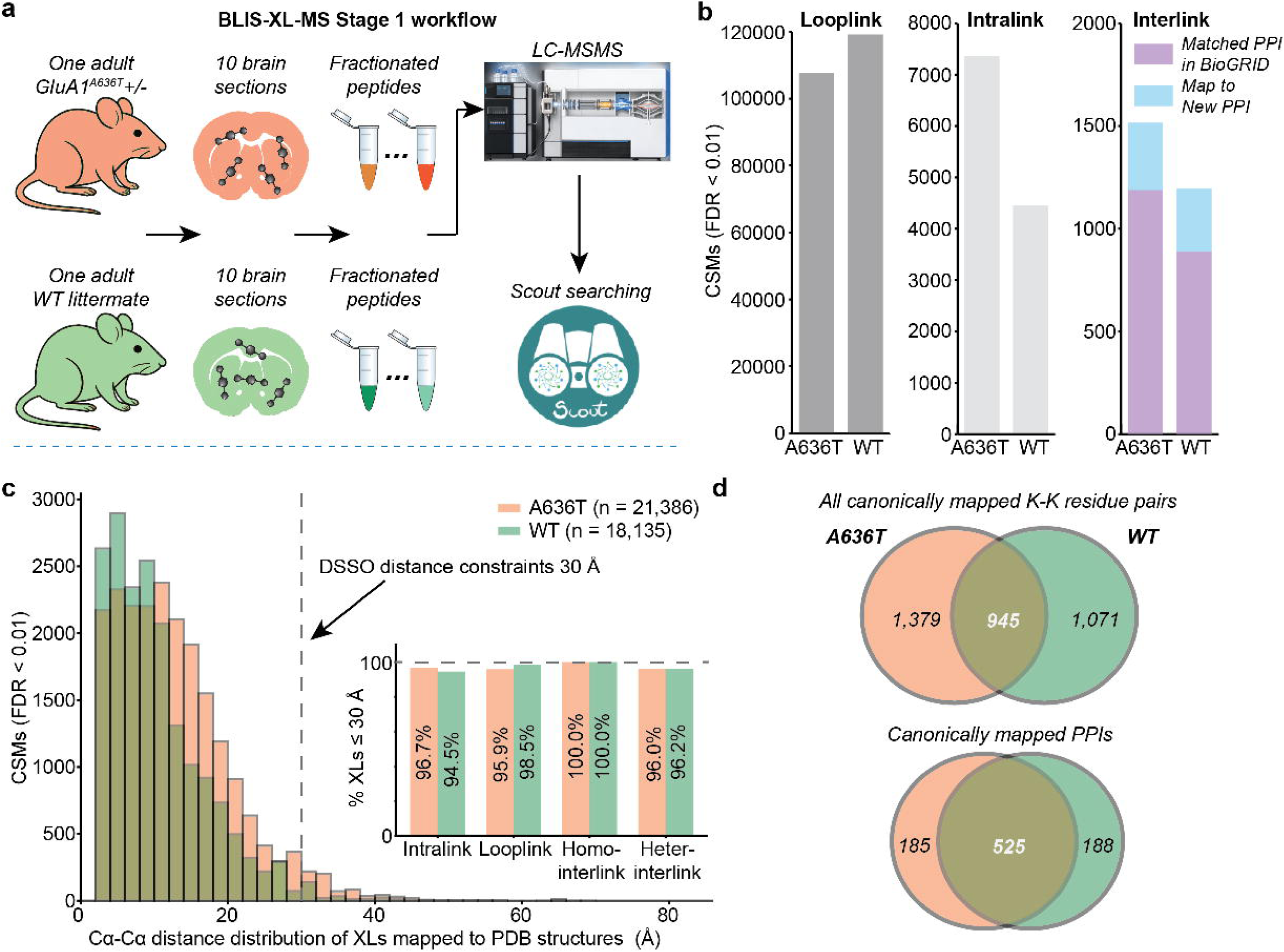
Quality assessment of BLIS-XL-MS crosslink identification and brain-proteome coverage. **a,** Sampling scheme for the BLIS-XL-MS quality-control dataset. One brain from a heterozygous GluA1^A636T^ mouse and one WT littermate was analyzed, using ten 50-µm sections per brain. Samples were processed through the BLIS-XL-MS stage 1 workflow, fractionated, and analyzed by LC-MS/MS. Raw files were searched with Scout, and crosslinked peptides were filtered at the CSM, residue-pair and PPI levels. Please also see **Extended Data Fig.2a**. **b,** Numbers of Scout-identified looplink, intralink and interlink CSMs passing CSM FDR < 0.01. A636T yielded 107,770 looplink, 7,356 intralink and 1,516 interlink CSMs; WT yielded 119,163, 4,450 and 1,196, respectively. For interlinks, stacked bars indicate whether the corresponding protein pair was previously reported in BioGRID or mapped to a PPI without prior BioGRID support. Of the interlink identifications, 1,187 of 1,516 (78.3%) in A636T and 888 of 1,196 (74.2%) in WT mapped to previously reported BioGRID interactions. **c,** Structural-compatibility assessment of BLIS-XL-MS restraints against experimentally determined protein structures in the PDB. Histograms show Cα-Cα distance distributions for crosslinks that could be mapped unambiguously to suitable PDB structures from the A636T and WT datasets. The vertical dashed line marks 30Å, used as the practical upper-distance bound for DSSO-derived restraints. Inset, percentage of structurally mapped crosslinks satisfying the ≤ 30Å criterion, stratified by link class. The corresponding percentages for A636T and WT were 96.7% and 94.5% for intralinks, 95.9% and 98.5% for looplinks, 100% and 100% for homomeric interlinks, and 96.0% and 96.2% for heteromeric interlinks, respectively. The audited datasets contained 21,386 unique crosslinks in A636T and 18,135 in WT. **d,** *Top*, all canonical K-K residue pairs mapped from all quality-filtered crosslinks: 1,379 were unique to A636T, 945 were shared between genotypes and 1,071 were unique to WT. *Bottom*, canonical PPIs: 185 were unique to A636T, 525 were shared and 188 were unique to WT. Values indicate protein counts after canonical mapping.

FDR threshold at the CSM, residue-pair, and PPI levels (**Fig. 2b**). Among canonically mapped XLs included in the structural assesment, 94.5-98.5% of looplinks and intralinks, 100% of evaluable homomeric interlinks and 96.0-96.2% of heteromeric interlinks satisfied a 30Å Cα-Cα DSSO restraint (**Fig. 2c & Extended Data Fig. 1b**). In addition, 78.3% of A636T and 74.2% of WT interlink-derived PPIs are supported by BioGRID (**Fig. 2b**, *right panel*). These orthogonal assesments support the structural and biological plausibility of the datasets.

Notably, looplink profiles showed substantially greater divergence between genotypes than intralink and interlink profiles (**Extended Data Fig. 2b**). We found that this discrepancy was largely driven by a mixture of low-quality crosslinks, which could disproportionately influence downstream remodeling analyses (**Extended Data Fig.2c-d**). Thus, BLIS 1.0 only retained high quality-filtered looplinks and combined them with intralinks for within-protein conformational- remodeling analysis (**Extended Data Fig.2e-f & Extended Data Fig.3)**.

To evaluate whether brain sectioning and Triton X-100 permeabilization improved DSSO accessibility to proteins across diverse structural and complex environments in brain tissue, we compared the crosslinked proteome with MS data from brain homogenates, which provide a reference for the broader set of brain proteins detectable by MS. We used our recently generated quantitative hippocampal proteome dataset from age-matched adult GluA1^A636T^ and WT mice, measured by 16-plex tandem mass tag (TMT)-MS (n = 8 A636T and n = 8 WT), as an orthogonal reference for this comparison (**Extended Data Fig.4a**)^16^.

To eliminate ambiguity arising from multiple isoforms, we first mapped proteins in both datasets to their canonical forms, ensuring that each crosslink was assigned to a single canonical protein or protein pair and that each TMT-labeled peptide was associated with a single canonical protein (**Extended Data Fig.1b**). As a result, Scout-identified crosslinks covered 34.2% of the mouse proteome in A636T tissue and 35.0% in WT tissue, compared with 24.5% coverage by TMT-MS (**Extended Data Fig.4b**). Moreover, BLIS XL proteomes showed a broadly similar distribution across major parent GO:CC categories, indicating that crosslinked proteins were recovered from most major subcellular compartments represented in the brain proteome (**Extended Data Fig.4c**). Together, these results suggest that brain sectioning and TX-100 permeabilization provided DSSO access to proteins across diverse intracellular compartments and structural environments, rather than restricting crosslinking to readily accessible protein populations.

Overall, BLIS-XL-MS workflow provides broad subcellular coverage without an obvious compartment-level or abundance-dependent bias that would systematically exclude major classes of proteins or protein complexes from downstream conformational-remodeling analyses.

#### BLIS 1.0 identifies within-protein conformational remodeling candiates linked to AMPAR trafficking and apoptosis in A636T brain

First, we used BLIS 1.0 to assess within-protein conformational remodeling using intralink and quality-filtered looplink CSMs (**Fig. 3**). Thirty proteins met Benjamini-Hochberg (BH)-FDR ≤ 0.05, including 17 with weighted JSD ≥ 0.20 that were designated high-effect conformational- remodeling candidates (**Fig. 3a & Extended Data Fig.5a-c**). STRING analysis of the 17 candidates prioritized Reactome terms related to AMPAR trafficking, mitochondrial function and cell death among the ten most significant categories (FDR < 0.01) (**Fig. 3b**). Related terms, except for ‘apoptosis’, were also recovered from TMT-MS-defined differentially abundant proteins. However, they ranked much lower within the substantially broader TMT-MS GO results (**Extended Data Fig.5d-f**). Notably, BLIS-XL-MS-implicated mitochondrial and cell-death signatures aligned with independently reported mitochondrial remodeling, oxidative stress, hippocampal degeneration and neuronal loss in two recent studies^16,17^.

**Fig. 3.**
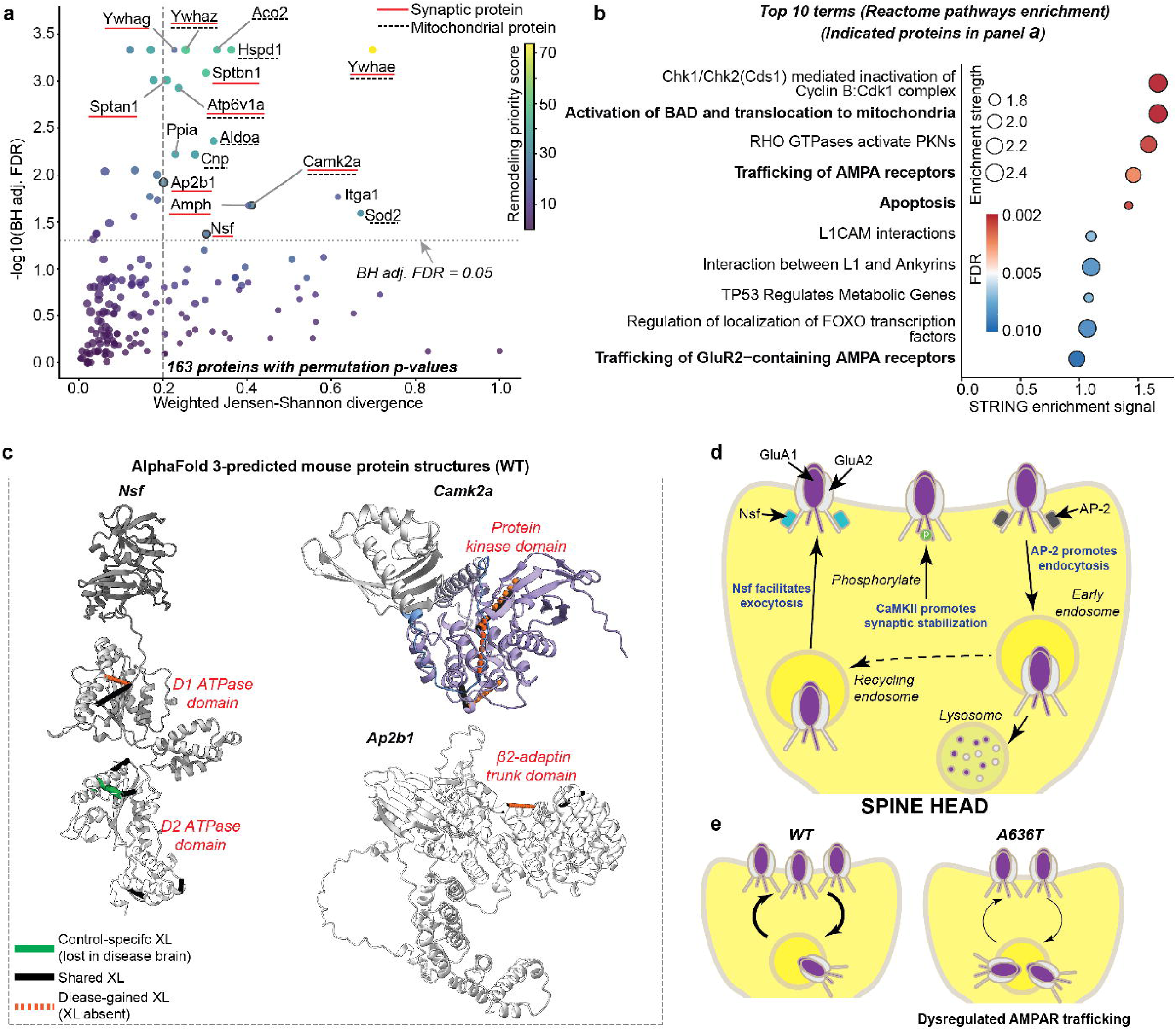
BLIS-XL-MS identifies within-protein conformational remodeling candidates linked to AMPAR trafficking in the A636T brain. **a,** Within-protein remodeling analysis using canonically mapped, quality-weighted intralink and quality-filtered looplink evidence. Each point represents one of the 163 proteins that met the evidence requirements for permutation testing. Point color indicates the BLIS remodeling-priority score (0–100), a technical prioritization metric integrating statistical support, structural-profile divergence, evidence depth, intralink contribution and gene-specific mapping confidence. Red underlines denote synaptic proteins, and black dashed underlines denote mitochondrial proteins. Please also see **Methods-**<u>Within-protein conformational-remodeling analysis using BLIS 1.0</u>. **b,** STRING Reactome pathway enrichment of the 17 high-effect conformational-remodeling candidates identified in **a**. The ten highest-ranked Reactome terms are shown. The x axis represents the STRING enrichment signal displayed in the analysis, point size represents enrichment strength and point color represents FDR. AMPAR-trafficking and cell-death-related terms are highlighted in bold. **c,** Structural localization of representative condition-associated crosslink patterns in three AMPAR-trafficking proteins prioritized by BLIS: NSF, CaMKIIα and AP2B1. Crosslinked lysine pairs were mapped onto AlphaFold 3-predicted WT mouse structures. For NSF and AP2B1, the remodeled protein is displayed in the structural context derived from the corresponding NSF and AP-2 models; CaMKIIα was modeled as a monomer because modeling of the complete oligomeric assembly exceeded the 5,000-token input limit of the AlphaFold 3 implementation used here. Remodelled residue-pair evidence localized to the D1 and D2 ATPase domains of NSF, the protein kinase domain of CaMKIIα and the β2-adaptin trunk region of AP2B1. Black lines denote crosslinks detected in both conditions, green lines denote crosslinks detected only in WT and orange dotted lines denote crosslinks detected only in A636T. Condition-restricted detection is used here to visualize the structural redistribution of XL evidence and should not be interpreted by itself as proof of biological gain or loss, because non-detection in XL-MS does not establish absence of the corresponding proximity. Model-confidence assessments are provided in **Extended Data Fig. 6.** **d,** Schematic illustrating the complementary functions of NSF, CaMKIIα and AP-2 in AMPAR trafficking. NSF contributes to receptor delivery and exocytosis, CaMKIIα-dependent phosphorylation promotes activity-dependent AMPAR stabilization at the synapse, and the AP-2 complex mediates clathrin-dependent receptor endocytosis. Internalized receptors can enter recycling pathways for return to the plasma membrane or be directed toward lysosomal degradation. GluA1- and GluA2-containing AMPARs are illustrated schematically. **e,** Working model for AMPAR-trafficking dysregulation in the A636T brain. In WT synapses, coordinated receptor delivery, stabilization, internalization and recycling maintain basal synaptic AMPAR occupancy. The BLIS-defined remodeling of NSF, CaMKIIα and AP2B1 is proposed to alter this balance in A636T, favoring a larger intracellular or recycling receptor pool and reduced basal surface/synaptic AMPAR occupancy. Arrow widths and receptor numbers are schematic and do not represent measured trafficking rates, stoichiometries or absolute receptor numbers.

The two AMPAR-trafficking terms were particularly notable. Three high-effect candidates, NSF, CaMKIIα and AP2B1, function at complementary stages of AMPAR trafficking, synaptic stabilization and endocytic removal^26–28^. Mapping remodeled residue pairs onto AlphaFold 3 models localized them to the D1 and D2 ATPase domains of NSF, the kinase domain of CaMKIIα and the β2-adaptin trunk region of AP2B1, suggesting coordinated reorganization of AMPAR-trafficking machinery (**Extended Data Fig. 6 & Fig. 3c**). This suggests a working model in which altered AMPAR delivery or stabilization shifts receptors toward intracellular or recycling pools, reducing basal synaptic occupancy and contributing to persistent silent synapses as previously reported in the GluA1^A636T^ hippocampus^16^. A larger pool of low-occupancy synapses could, in turn, contribute to the greater potentiation observed in A636T than in WT following strong stimulation, through activity-dependent AMPAR recruitment (**Fig. 3d**).

Therefore, BLIS-XL-MS reveals a previously unrecognized mechanistic link between the A636T gating defect and synaptic dysfunction, implicating coordinated remodeling of AMPAR trafficking machinery as a molecular basis for persistent synaptic immaturity and aberrant plasticity in the adult A636T brain^16,17^.

#### BLIS 1.0 reveals reveal Complex V remodeling in the A636T brain

We next analyzed interlinks to identify protein-complex remodeling in the GluA1^A636T^ brain (**Fig. 4**). Proteins were represented as nodes and interlink-derived PPIs as edges in a condition-blind consensus network. At the overall network level, the strongest A636T-associated PPIs were clustered within one major interaction network, whereas weaker interactions were more scattered across smaller peripheral groups (**Fig. 4a-c**). This pattern suggests concentrated remodeling of core protein assemblies together with broader changes in weaker interactions.

**Fig. 4.**
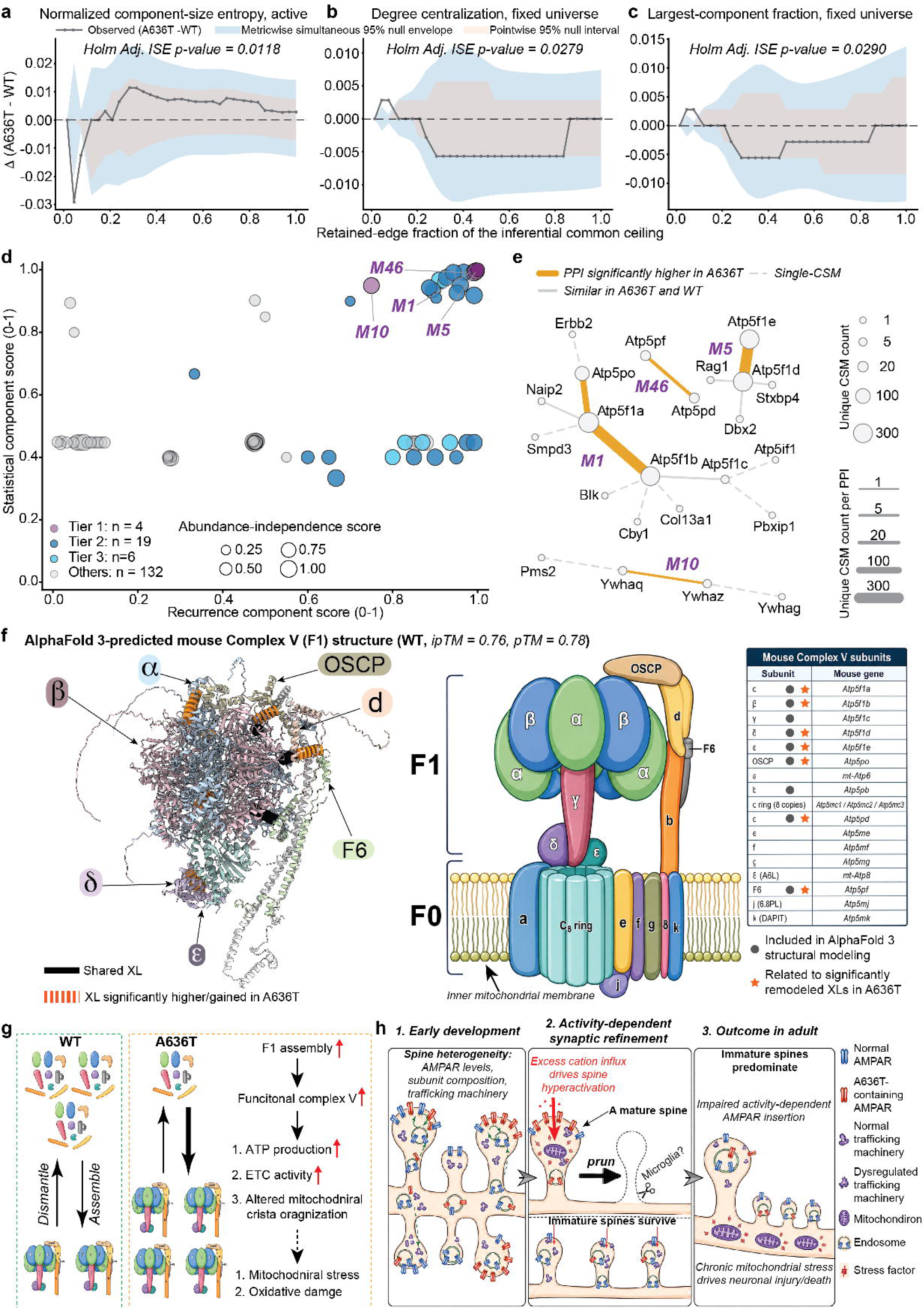
BLIS-XL-MS identifies protein-complex remodeling candidates centered on mitochondrial Complex V in the A636T brain. **a-c,** Forest-aware, equal-edge-count comparison of global PPI-network organization between A636T and WT across the retained-edge filtration. At each point, the same number of top-ranked positive heteromeric PPI edges was retained for each condition; the x axis indicates the retained- edge count as a fraction of the inferential common ceiling. Curves show the observed difference, A636T minus WT. Shaded regions indicate permutation-derived null intervals, including the metric-wise simultaneous 95% null envelope and pointwise 95% null interval. Statistical significance was assessed over the complete filtration using an integrated-squared-error (ISE) curve statistic, with Holm correction across the tested topology metrics. Because the retained PPI networks were acyclic forests, the analysis emphasized component organization and centralization rather than triangle-dependent measures such as clustering or transitivity. **a,** Difference in normalized component-size entropy calculated among active, non-isolated proteins (Holm-adjusted ISE p-value = 0.0118). **b,** Difference in degree centralization calculated over the fixed protein universe, thereby retaining changes in protein participation as part of the network phenotype (Holm-adjusted ISE p-value = 0.0279). **c,** Difference in the fraction of the fixed protein universe contained within the largest connected component (Holm-adjusted ISE p-value = 0.0290). Collectively, the filtration profiles indicate altered organization of the A636T interaction network, with the strongest A636T-supported PPIs preferentially concentrating within a dominant component and lower-ranked interactions showing greater fragmentation into peripheral components. Please also see **Methods-**<u>Interlink-based protein-complex remodeling analysis using BLIS 1.0</u> **d,** Integrated technical-evidence prioritization of PPI remodeling objects (*n* = 161), comprising dyadic PPIs, small complexes and condition-blind network modules. Each point represents one PPI object and is positioned according to its recurrence-component score and statistical- component score; point size represents its abundance-independence score. Component scores are scaled from 0 to 1 within object class, with larger values indicating stronger support in the corresponding evidence dimension. The abundance-independence component summarizes interaction remodeling remaining after accounting for protein-abundance changes measured by TMT-MS. Colors denote technical-evidence tiers: Tier 1, statistical and recurrent support (*n* = 4); Tier 2, strong technical candidates (*n* = 19); Tier 3, recurrent descriptive candidates (*n* = 6); and other objects (*n* = 132), comprising exploratory and sparse/singleton objects. The four Tier-1 objects, M1, M5, M46 and M10, are labelled. **e,** Interaction maps of the four Tier-1 PPI-remodeling objects. M1, M5 and M46 converge on mitochondrial ATP synthase (Complex V), whereas M10 is centered on 14-3-3 proteins. Orange edges denote PPIs with significantly greater BLIS interlink evidence in A636T, solid grey edges denote PPIs with comparable evidence between A636T and WT, and dashed grey edges indicate single-CSM-supported interactions retained for network context. Node size represents unique CSM support, and edge width scales with the number of unique CSMs supporting each PPI. **f,** Structural localization of Complex V remodeling. *Left*, AlphaFold 3-predicted WT mouse F1- containing Complex V assembly used to provide structural context for the BLIS interlink evidence (ipTM = 0.76, pTM = 0.78). Subunits are shown in distinct colors and representative remodeled interfaces are indicated. Black lines denote XLs shared between A636T and WT, whereas orange marks denote XL evidence significantly increased in, or detected selectively in, A636T. *Right*, schematic organization of mammalian Complex V, comprising the soluble F1 catalytic sector and membrane-embedded F0 sector. The accompanying table lists mouse genes corresponding to individual Complex V subunits; filled grey circles identify subunits included in the AlphaFold 3 model and orange stars identify subunits associated with significantly remodeled A636T XL evidence. The remodeled interfaces span the F1 α/β catalytic head, central-stalk and peripheral- stalk regions, placing the independently prioritized BLIS interactions within multiple structural elements required for assembly and function of ATP synthase. Condition-restricted XL detections are interpreted as remodeling evidence rather than proof that the corresponding molecular contact is biologically absent in the other condition. Model-confidence assessment is shown in **Extended Data Fig. 7**. **g,** Working model for Complex V remodeling in A636T brain. In WT, ATP synthase subunits and subcomplexes are depicted as dynamically partitioning between less assembled and fully assembled states. The coordinated increase in interlink evidence at several Complex V interfaces in A636T is consistent with a redistribution toward more assembled F1/Complex V states. Such remodeling could support increased ATP-synthase activity and ATP production and may accompany altered electron-transport-chain activity and mitochondrial crista organization. Sustained enhancement of mitochondrial energetic activity is proposed to increase susceptibility to mitochondrial stress and oxidative damage. Arrows and relative complex abundances are schematic and do not represent direct measurements of Complex V assembly kinetics, ATP- production rates or stoichiometry. **h,** Integrated working model linking the primary A636T AMPAR gating defect with synaptic- trafficking and mitochondrial remodeling. <u>1, Early development:</u> developing excitatory synapses exhibit heterogeneity in AMPAR abundance and subunit composition, trafficking machinery and activity. <u>2, Activity-dependent synaptic refinement:</u> enhanced cation influx through gain-of- function A636T-containing AMPARs is proposed to preferentially hyperactivate highly active developing synapses, promoting their elimination during activity-dependent refinement, whereas less active or immature synapses preferentially persist. A potential contribution of microglia- mediated pruning is indicated as a hypothesis. <u>3, Adult outcome:</u> remodeling of NSF, CaMKIIα and AP2B1-dependent trafficking pathways is proposed to impair normal AMPAR delivery and activity-dependent synaptic stabilization, favoring persistent low-AMPAR or immature synaptic states. In parallel, increased energetic demand and Complex V reorganization may contribute to chronic mitochondrial and oxidative stress, ultimately increasing susceptibility to neuronal injury or death. Symbols denote normal and A636T-containing AMPARs, normal and remodeled trafficking machinery, mitochondria, endosomes and cellular-stress signals.

We then examined individual edges and condition-blind consensus modules (**Fig. 4d-e & Extended Data Fig.7a-b**). The top four remodeled modules were the ATP synthase-centered modules M1, M5 and M46, and the 14-3-3-centered module M10. Thus, three modules converged on mitochondrial Complex V, spanning the F1 α/β head, central stalk and peripheral-stalk interfaces (**Extended Data Fig.7c & Fig. 4f**). These findings suggest a shift in the distribution of Complex V assembly states toward more assembled ATP synthase in the A636T brain (**Fig. 4g**). Such a shift could enhance Complex V activity and ATP production, potentially contributing to mitochondrial remodeling and increased metabolic stress^29–32^.

Integrating the within-protein conformational and protein-complex remodeling results, our analyses support a working model in which gain-of-function A636T-containing AMPARs preferentially destabilize highly active developing synapses, while remodeling of NSF, CaMKIIα, and AP2B1 may reflect altered AMPAR trafficking and reduced basal synaptic AMPAR occupancy, thereby favoring persistent silent synapses, as demonstrated by previous functional studies (**Fig. 4h**)^33–36^. In parallel, sustained synaptic remodeling may increase energetic demand, with Complex V reorganization supporting this demand while increasing susceptibility to redox stress and mitochondrial dysfunction. Together, these coordinated synaptic-trafficking and mitochondrial alterations provide a structural and interaction-level framework linking the A636T variant to persistent synaptic immaturity, aberrant plasticity and progressive neuronal injury^16,17^.

## Discussion

The development of the BLIS-XL-MS experimental workflow together with the BLIS 1.0 computational package enables proteome-scale profiling of within-protein conformational and protein-complex remodeling with quantitative statistical support. By providing structural and interaction-level information that complements conventional abundance-based proteomics, BLIS- XL-MS can prioritize candidates for targeted validation, generate structure-guided mechanistic hypotheses, and identify candidate interfaces or conformationally sensitive regions for drug development. More broadly, the framework could be applied to investigate disease-associated structural remodeling, PTM-dependent conformational changes and drug-induced or ligand-binding structural responses, supporting diverse applications across biology, biochemistry and translational research.

Notably, the fixation and secondary-crosslinking strategy does not eliminate *ex vivo* protein conformational remodeling (**Extended Data Fig. 8a**). Rather, it shifts the principal state- capture window from the prolonged secondary-crosslinking reaction to the preceding fixation step (**Extended Data Fig. 8b**). Rapid delivery of PFA, particularly by perfusion, can substantially shorten this vulnerable interval, and fixation at low temperature (e.g., 4°C) can further limit post-dissection molecular dynamics compared with prolonged crosslinking of viable tissue (e.g., 37°C). However, many specimens of practical interest, particularly human brain tissues, are obtained by dissection followed by immersion fixation. We therefore evaluated BLIS-XL-MS using post-fixed mouse brains, despite recognizing that perfusion fixation should provide better molecular-state preservation, to test whether immersion-fixed tissue remains suitable for comparative structural proteomics. The high proportion of crosslinks satisfying the 30Å Cα-Cα restraint (> 94.5%), together with substantial agreement of interlink-derived PPIs with BioGRID (> 72.2%), indicates no obvious large-scale structural or interaction distortion. These findings suggest that the BLIS- XL-MS workflow is compatible with post-fixed brain tissue, including immersion-fixed postmortem human brain samples. Although we have not yet tested BLIS-XL-MS in other tissue types, such as liver or heart, the underlying principles suggest that the workflow may be broadly applicable to other tissue types.

For the within-protein and protein-complex remodeling events identified by BLIS 1.0, the results should not be interpreted as NSF, AP-2, CaMKIIα, or Complex V adopting one discrete structure in A636T brain and a different single structure in WT brain. Proteins and protein complexes are not restricted to single, rigid structures. Many populate ensembles of interconverting conformations, and interactions with binding partners can preferentially stabilize particular states^37,38^. In neurons, developmental maturation regulates the composition and higher-order assembly of receptor complexes, while activity-dependent biochemical changes can promote persistent organization of postsynaptic proteins^39–41^. Assembly-dependent quality control provides an additional mechanism for regulating molecular persistence, including the selective removal of certain unassembled subunits. Together, our findings support a framework in which cellular context shapes the relative populations and persistence of protein conformations and assemblies, rather than universally selecting a single optimal structure.

Proteome-scale quantitative comparison of XL-MS datasets remains analytically challenging. Multiplexed strategies such as TMT labeling can enable quantitative XL-MS^15,42^. However, they add peptide-derivatization steps and increase search complexity because crosslinked peptides already contain multiple variable modifications, cleavage states and linked- peptide combinations. BLIS 1.0 instead operates directly on label-free XL-MS datasets and does not require additional chemical labeling or specialized acquisition. Therefore, BLIS 1.0 analysis requires only modest computational resources. Thus, sample generation and MS analysis can remain compatible with standard XL-MS workflows available in many proteomics laboratories, while BLIS 1.0 provides the downstream statistical framework for comparative structural and interaction-level analysis.

Notably, BLIS 1.0 is currently designed for pairwise statistical comparisons. Although it yielded biologically coherent findings in this proof-of-concept study, the framework does not fully leverage low-support evidence, including low-abundance singleton crosslinks, and its empirical permutation statistics primarily quantify technical rather than population-level biological variation. We are therefore developing BLIS 2.0 to integrate evidence across multiple matched biological pairs, improve crosslink coverage and statistical power, and combine within-pair permutation evidence with replicate-level biological inference.

Together, BLIS-XL-MS provides an accessible experimental and computational framework for extracting structural and interaction-level information from limited or irreplaceable brain specimens, extending conventional proteomics beyond protein abundance to reveal how proteins and protein complexes are organized and remodeled in tissue.

## Methods

### Animals

All procedures were approved by Northwestern University’s Animal Care and Use Committee (IS00020323) in compliance with US National Institutes of Health standards. Recently, we generated the *Gria1* A636T knock-in mouse line. Heterozygous A636T mice were used as breeders to generate 3-month-old A636T and WT littermate controls for the experiments.

### In-tissue crosslinking

Mice were decapitated, and brains were rapidly removed and immersion-fixed overnight at 4 °C in PBS containing 4% PFA (pH 7.4). Coronal brain sections (50 µm thick) were prepared using a Leica vibratome and washed six times with PBS (MEDIATECH INC CA, Cat# 21-040-CV) to remove residual PFA. Sections were permeabilized in 0.1% Triton X-100 (Sigma-Aldrich, Cat# X100) in PBS for 30 min at room temperature with gentle rotation, followed by three 5-min washes in PBS.

Ten brain sections per genotype were used for crosslinking. Every five brain sections were incubated in 1 ml of freshly prepared 2 mM DSSO (Fisher Scientific, Cat# A33545) in PBS (pH 7.4) for 30 min at room temperature with rotation. Sections were then washed three times with PBS for 5 min each to remove hydrolyzed DSSO and incubated with a second freshly prepared 1 ml aliquot of 2 mM DSSO in PBS (pH 7.4) for an additional 30 min under the same conditions. After one PBS wash, residual reactive DSSO was quenched with 50 mM Tris-base for 15 min at room temperature with rotation.

### MS sample preparation

Brain sections were collected into a PFA-reversal lysis buffer containing 500 mM Tris-base, 150 mM NaCl, 1% Triton X-100, 2% SDS, 50 mM 1,4-dithiothreitol (DTT), and 1× protease and phosphatase inhibitor cocktail (Thermo Fisher Scientific, Cat# 78443), pH 8.0. Samples were then incubated at 95 °C for 1 h with shaking at 1,000 rpm. After cooling to room temperature, samples were homogenized using a Precellys 24 homogenizer (Bertin Technologies, Cat# P000669- PR240-A) with four 20-second pulses at 5,000 rpm. Samples were denatured in 8 M urea (Thermo Fisher Scientific, Cat# 29700) for 30 min at room temperature. Iodoacetamide (IAA, Sigma-Aldrich, Cat# I1149) was then added to a final concentration of 15 mM, and samples were incubated for 20 min at room temperature in the dark. Excess IAA was quenched with DTT for 15 min. Samples were subsequently cleaned up using SP3 beads and resuspended in 2 M urea in 100 mM ammonium bicarbonate (Fluka, Cat. no. 09830). Proteins were digested with Lys-C (1:50 enzyme-to-protein ratio, Thermo Fisher Scientific, Cat# 90307) for 3 h at 37 °C, followed by addition of trypsin (1:100, Promega, Cat# V5280) and overnight digestion at 37 °C with shaking at 1,000 rpm. The next day, reaction was quenched by adding 1% trifluoroacetic acid (TFA, Fisher Scientific, Cat# O4902-100). Samples were desalted using Peptide Desalting Spin Columns (Thermo Fisher Scientific, Cat# 89852). All samples were vacuum centrifuged to dry.

The peptides were fractionated by strong cation exchange (SCX) chromatography using a HyperSep SCX SPE column (Thermo Fisher Scientific, Cat# 60108-420) with sequential elution at 20, 50, 100, 200, 350, 500, 750, 1,000, 1,500, and 2,000 mM ammonium acetate (Sigma-Aldrich, Cat# 09689). The SCX fractions were desalted using Peptide Desalting Spin Columns. The 50, 100, 200, 350, 500, 750, and 1,000 mM SCX fractions were further fractionated by high-pH reversed-phase chromatography using the High pH Reversed-Phase Peptide Fractionation Kit (Thermo Fisher Scientific, Cat# 84868). Peptides were sequentially eluted into nine fractions using 10.0%, 12.5%, 15.0%, 17.5%, 20.0%, 22.5%, 25.0%, 50%, and 80% acetonitrile (ACN) in 0.1% triethylamine. The resulting high-pH reversed-phase fractions were loaded directly into the autosampler for MS analysis without additional desalting.

### LC-MS/MS analysis

Samples were loaded using an autosampler with a Thermo Vanquish Neo UHPLC system onto a PepMap Neo Trap Cartridge (Thermo Fisher Scientific, 174500; diameter, 300 µm; length, 5 mm; particle size, 5 µm; pore size, 100 Å; stationary phase, C18) coupled to a nanoViper analytical column (Thermo Fisher Scientific, 164570; diameter, 0.075 mm; length, 500 mm; particle size, 3 µm; pore size, 100 Å; stationary phase, C18) with a stainless-steel emitter tip assembled on the Nanospray Flex Ion Source with a spray voltage of 2,200 V. An Orbitrap Ascend (Thermo Fisher Scientific) was used to acquire all the MS spectral data. Buffer A contained 99.9% H2O and 0.1% formic acid, and buffer B contained 80.0% acetonitrile, 19.9% H2O with 0.1% formic acid. For each high-pH fraction, approximately 1 μg of peptides was loaded for LC-MS/MS analysis. The total chromatographic run time was 3 h. For the first fraction generated by high-pH reversed-phase fractionation, using the following gradient profile: 2-20% B over 154 min, 20-55% B over 16 min, 55-80% B over 1 min, 80-8.8% B over 1 min, held at 8.8% B for 2 min, followed by 99% B for 6 min. For the rest high pH fractions: 8.8-37.5% B over 114 min, 37.5-55% B over 56 min, 55-80% B over 1 min, 80-8.8% B over 1 min, held at 8.8% B for 2 min, followed by 99% B for 6 min. The flow rate is 350 nL/min at 50 °C.

Instrument parameters were set as follows. MS1 spectra were acquired in the Orbitrap at a resolution of 60,000 over an m/z range of 380-1,600, with the AGC target set to ‘Standard’ (absolute AGC target, 4.0 × 10E5) and the maximum injection time set to ‘Auto’. The precursor intensity threshold was set to 2.5 × 10E4. The default precursor charge state was set to 4+, with precursor selection restricted to charge states 3+ to 8+, and dynamic exclusion was set to 60 s. Precursors were fragmented by stepped higher-energy collisional dissociation (HCD) using normalized collision energies of 21%, 27%, and 33%. MS2 spectra were acquired in the Orbitrap at a resolution of 30,000 over an m/z range of 145-2,000, with the AGC target set to 200% and the maximum injection time set to 100 ms.

### Scout search

Raw XL-MS files were individually searched using Scout (version 2.1.1 for Linux) against the Mus musculus UniProt reference proteome database (UP000000589; database downloaded on 16 October 2025). Searches were performed in batch automation mode under Ubuntu 22.04 Linux using Windows Subsystem for Linux 2 (WSL2), with parallel processing on a workstation providing 24 CPU cores. Search parameters included a minimum peptide length of 6 amino acids, a maximum peptide length of 60 amino acids, up to 3 missed cleavages and a maximum of 2 variable modifications per peptide. Carbamidomethylation of cysteine (+57.02146 Da) was specified as a fixed modification and methionine oxidation (+15.9949 Da) as a variable modification. DSSO was specified as the MS-cleavable crosslinker using a full crosslinker mass of 158.00376533 Da and diagnostic short- and long-arm fragment masses of 54.01056468 and 85.98263585 Da, respectively. Only K is permitted as crosslinking site. CSMs were filtered using Scout with FDR thresholds of 1% at the CSM, residue-pair and PPI levels. Scout-filtered CSM, looplink, residue-pair and PPI reports were exported for subsequent BLIS 1.0 analyses.

### Within-protein conformational-remodeling analysis using BLIS 1.0

BLIS 1.0 was developed for the analysis of XL-MS datasets generated using lysine-reactive crosslinkers that predominantly yield K-K residue pairs (**Fig. 1b**, *left*). It is configured by default to use Scout output files as input for downstream analysis. Within-protein conformational remodeling was analyzed using BLIS 1.0 directly from the CSMs and looplink reports exported by Scout. BLIS 1.0 was configured for K-K residue-pair evidence, and each row of a Scout output file was treated as one CSM event. Following the Scout FDR filtering described above, target intralink CSMs with Scout scores ≥ 0.00 and target looplink CSMs with Scout scores ≥ 0.20 were retained. Interlinks were excluded from this analysis and were analyzed separately in the PPI-remodeling part below. All feature construction, evidence weighting, normalization and statistical testing were performed directly in BLIS 1.0. No pre-normalized abundance estimates or differential-statistical results from another XL-MS analysis pipeline were imported.

In the equations below, *i* indexes CSMs, *g* indexes genes or proteins, and *λ* ∈ ℒ_g_ indexes canonical K-K residue-pair features of protein *g*. Each feature was defined as *λ* = (*τ_λ_*, *κ_λ_*_1_, *κ_λ_*_2_), where *τ_λ_* ∈ intralink, looplink denotes the link class and *κ_λ_*_1_ < *κ_λ_*_2_ are the positions of the two linked lysines in the selected canonical protein sequence. The symbols *r*, *s*, *c* ∈ *A*, *W* and *b* = 0, …, *B* index an LC-MS/MS run, an SCX salt stratum, condition and a restricted label assignment, respectively, with *A* denoting A636T, *W* denoting WT and *b* = 0 denoting the observed assignment.

For looplinks, each complete two-residue localization hypothesis reported in the Scout ‘ReactionSites (%)’ field was treated as one pair-level hypothesis. Duplicate pair-level hypotheses were merged by summing their reported percentages, and positive, finite reaction-site percentages were normalized to sum to one and treated as fractional localization weights. For intralinks, reaction-site percentages were normalized separately for the α- and β-peptides, and the fractional weight of each possible α-β site combination was calculated as the product of the corresponding peptide-specific weights. This product was used as a factorized evidence- allocation rule and was not interpreted as a calibrated joint posterior probability. Localization weights were normalized before the K-K restriction was applied. Consequently, evidence assigned by Scout to a non-K-K or otherwise inadmissible localization was not redistributed to the remaining K-K alternatives.

Scout reaction-site positions were first converted to source-protein coordinates using the corresponding peptide- and protein-position anchors. Source accessions were then mapped to a gene-specific canonical coordinate system using the same FASTA database employed for the Scout search. Canonical accessions were selected using a reviewed-primary-longest hierarchy, which preferentially selected a reviewed, non-fragment, unsuffixed primary accession before considering protein length; prespecified canonical-accession overrides were applied where required. Alternative isoforms were globally aligned to the selected canonical sequence using match, mismatch, gap-opening and gap-extension scores of 2, −1, −8 and −0.5, respectively, with unpenalized terminal gaps. A source-to-canonical mapping was accepted only when alignment identity was ≥ 0.90 and the source residue mapped to the same canonical position across all examined equally optimal alignments. Up to 50 equally optimal alignments were examined. Mappings for which positional consensus could not be verified were left unresolved. Retained canonical positions were required to contain lysine, and mappings were removed when the two endpoints collapsed onto the same canonical residue. For intralinks, the two endpoints were required to map to the same canonical accession, and cross-accession intralinks were not permitted.

Let *w*^loc^ denote the normalized localization weight of localization hypothesis ℎ for CSM *i*. For each hypothesis, V*_i_*_ℎ_ denotes the set of distinct valid canonical outcomes (*g*, *λ*), and U^coord^ denotes the set of distinct unresolved canonical-coordinate outcomes. The set ℋ^∗^ contains localization hypotheses having at least one valid canonical outcome. Under the non-renormalizing unresolved-outcome policy, a valid outcome received the corresponding localization weight divided by the total number of valid and unresolved outcomes. Cross-gene ambiguous CSMs were retained for auxiliary reporting but excluded from the primary within-protein profile. Thus, the fractional primary-profile contribution of CSM *i* to feature *λ* of protein *g* was:

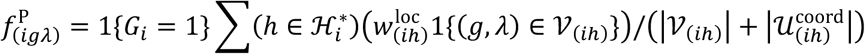

where *G_i_* is the number of source-level gene-mapping contexts represented by CSM *i*, including an additional unresolved context when an unannotated source mapping was present. This fractional allocation ensured that one Scout CSM contributed no more than one CSM-equivalent of evidence across all retained canonical residue-pair assignments. Evidence associated with inadmissible localizations or unresolved mappings remained unassigned rather than being transferred to the surviving assignments.

Each fractional CSM assignment was subsequently weighted according to Scout score, precursor-mass accuracy, link class and canonical sequence separation. Score and precursor- error weights were calibrated separately for intralinks and looplinks after pooling retained CSMs from both conditions, thereby avoiding condition-specific evidence scales. Within each link class, Scout scores and absolute precursor errors were linearly scaled between their pooled 5th and 95th percentiles and clipped to the interval [0,1]. The precursor-error measure was the larger available absolute ppm error of the α- and β-peptide assignments. The resulting score and precursor-error weights were 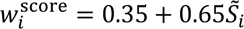 and 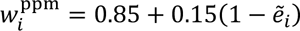 respectively, where *S̃_i_* and *ẽ_i_* are the scaled Scout score and absolute precursor error. If an entire link class contained no finite observations for a metric, its corresponding final weight was set to 1.0. Otherwise, an individual missing value or a degenerate 5th-95th percentile interval was assigned a scaled value of 0.5.

Fixed link-class weights of 1.00 and 0.60 were applied to intralinks and looplinks, respectively. Canonical sequence separation *d_λ_*, the separation weight 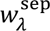, and the final quality-weighted primary-profile contribution was:

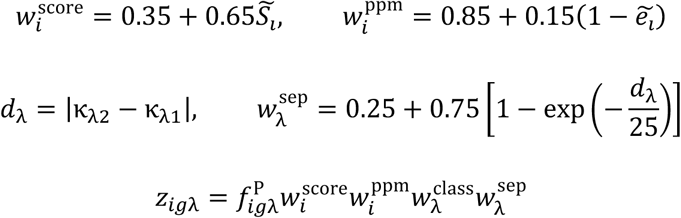

Weighted evidence was then aggregated separately for each protein, canonical residue-pair feature and LC-MS/MS run. Let ℐ_g*rλ*_ be the set of CSM assignments from run *r* that contributed to feature *λ* of protein *g*. The effective fractional CSM mass *E*_g*rλ*_, quality-weighted mass *Q*_g*rλ*_, mean reliability *w̄*_g_*_rλ_*, peptide-form diversity factor *D*_g*rλ*_ and final run-level support *x*_g*rλ*_ were calculated as:

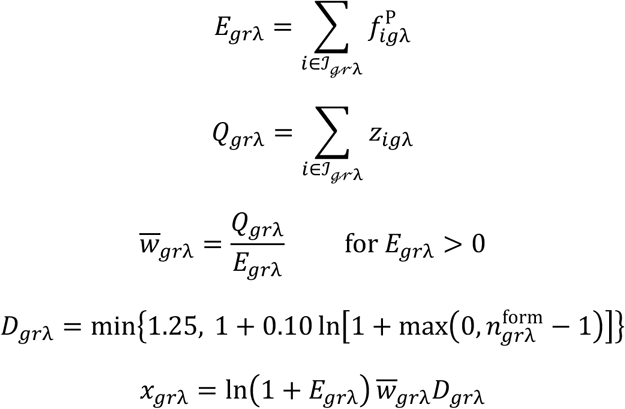

Here, 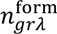 is the number of distinct peptide forms supporting the residue-pair feature, with each form defined by the α- and β-peptide sequences and their reported modifications. For looplinks, the absent β-peptide fields were represented by empty values in the peptide-form signature. When *E*_g*rλ*_ = 0, *w̄* _g*rλ*_ and *x*_g*rλ*_ were set to zero. The l n(1 + *E*) transformation imposed diminishing returns on repeated observations within one raw file, preventing heavily sampled CSMs in a single run from overwhelming evidence recurring across independent runs or fractions.

LC-MS/MS runs were grouped into six SCX salt strata corresponding to fractions eluted with 50, 100, 200, 350, 500 and 750 mM ammonium acetate^43^. Only strata containing at least two LC-MS/MS runs from each condition were retained, and all downstream evidence, eligibility and statistical calculations were restricted to runs in these retained strata. Let S denote the retained stratum set, ℛ*_cs_* the runs assigned to condition *c*in stratum *s*, and *n_cs_* =∣ ℛ*_cs_* ∣. Run-level supports were first averaged within each condition and salt stratum and then averaged across strata:

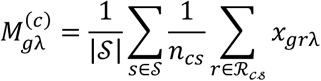

This assigned equal weight to each retained SCX stratum irrespective of the number of LC-MS/MS runs that it contained. Within a condition-stratum combination, each run received equal weight. A protein absent from a valid run contributed zero support in that run. The run was not removed from the averaging denominator on a protein-by-protein basis.

Proteins were subjected to permutation testing only when they contained at least five primary-profile fractional CSM equivalents and at least one gene-specific fractional CSM equivalent in each condition, were detected in at least two retained LC-MS/MS runs per condition, and contained at least three combined canonical residue-pair features contributing ≥ 1% of the unsmoothed within-protein profile in at least one condition. The 1% criterion was used only to count supported features for statistical testability. Features contributing < 1% were not removed and remained included in the subsequent Jensen-Shannon divergence (JSD) calculation.

For each eligible protein, ℒ_g_ was defined as the union of its retained canonical residue-pair features across both conditions, and *L*_g_ =∣ ℒ_g_ ∣. The SCX-balanced feature masses were normalized to unit sum separately within each protein and condition. A total pseudocount of *ε* = 10^−8^, divided equally among all features, was added as a numerical safeguard:

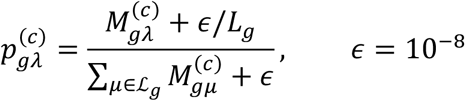

Dissimilarity between the A636T and WT profiles was quantified using base-2 JSD. Defining the midpoint profile as 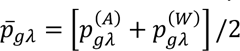, the observed protein-level statistic was:

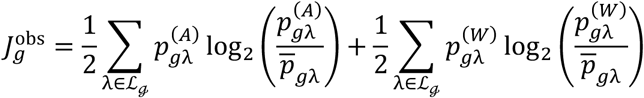

With base-2 logarithms, 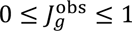, with zero indicating identical profiles and one indicating profiles with completely non-overlapping support. The term “weighted JSD” refers to conventional JSD applied to the preceding quality-weighted, run-aggregated and SCX-balanced profiles. Thus, the mathematical definition of JSD itself was not modified.

Statistical significance was evaluated by restricted whole-run label permutation. The run-by-feature support matrix *x*_g*r*_*_λ_* was calculated before condition-level averaging and was held fixed throughout the permutation procedure. All residue-pair values belonging to one LC-MS/MS run were therefore reassigned together. Within each retained SCX stratum, condition labels were randomly reassigned while preserving the observed numbers of A636T and WT runs. For permutation *b*, let 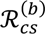 denote the runs assigned to condition *c* in stratum *s*, where 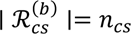 The permuted condition-level feature masses were:

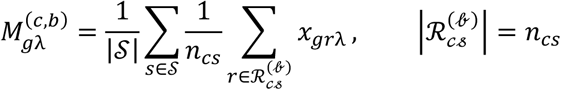

The within-protein profiles and JSD were recalculated for each permuted labeling using equations above. For *B* restricted permutations, the one-sided empirical *P* value was calculated with the plus-one correction:

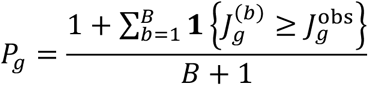

The reported analysis used *B* = 49,999 permutations and random seed 659660. A numerical tolerance of 10^−12^ was used when determining whether a permuted JSD tied or exceeded the observed statistic. Protein eligibility was determined once under the observed run labels and was not recalculated for individual permutations. Empirical *p* −values were adjusted across all eligible proteins using the Benjamini-Hochberg procedure. Proteins with BH-FDR ≤ 0.05 were considered to show significant within-dataset evidence of conformational remodeling, and significant proteins with weighted JSD ≥ 0.20 were designated high-effect candidates.

### Interlink-based protein-complex remodeling analysis using BLIS 1.0

Protein complex remodeling was analyzed in BLIS 1.0 using interlink CSMs exported from Scout v.2.1.1, also after filtering at a 1% FDR at the CSM, residue-pair and PPI levels (**Fig. 1b**, *right*). The downstream analysis was conditional on the resulting Scout-identified feature set. Intralinks and looplinks were excluded from the PPI analysis and analyzed separately in the within-protein conformational-remodeling workflow above. BLIS 1.0 used trimmed mean of M-values with singleton pairing (TMMwsp) normalization followed by log2-transformed counts per million (logCPM) transformation for run-aware edge modeling, together with within-run edge compositions for network-level comparisons^44–49^. Notably, homomeric evidence was retained for edge analysis, whereas self-loops were excluded from topology and module-connectivity calculations^50–52^.

### Interlink evidence processing and construction of protein-pair edges

Each retained Scout row was treated as one CSM event. Gene annotations for the two crosslinked peptides were parsed independently, and all possible α-β combinations were converted to canonical, unordered gene pairs. Duplicate pair hypotheses arising within the same CSM were collapsed. Let ℰ*_i_* denote the set of distinct candidate gene-pair edges for CSM *i*, let *e* = *g*, ℎ denote an unordered gene-pair edge and let *r_i_* denote the LC-MS/MS run containing CSM *i*. Under the primary unambiguous assignment rule:

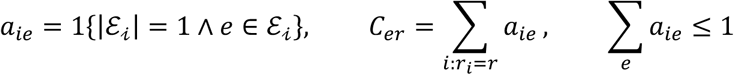

Thus, a CSM compatible with exactly one gene pair contributed one unit of evidence to that edge, whereas a CSM compatible with more than one gene pair contributed no mass to the primary edge-level matrix. Ambiguous CSMs were retained in the mapping audit. A secondary mapping-sensitivity analysis fractionally distributed one unit of CSM mass among all candidate gene-pair edges, but this analysis was not used for the primary results.

Both heteromeric gene pairs (heter-) and same-gene (homo-) interlinks were retained in the 213-edge family used for edge-level and occupancy analyses. Homo-interlinks were not interpreted as unequivocal homomeric contacts because the observed peptide pair may not distinguish intermolecular homomeric proximity from alternative intramolecular or proteoform configurations. Conventional network topology and module analyses excluded same-gene self-loops and were therefore restricted to 197 heteromeric edges spanning a fixed universe of 357 proteins. No edge was excluded solely because it was supported by a single CSM. Instead, CSM count, recurrence across runs and recurrence across SCX strata were retained as evidence-strength annotations.

### Run-level normalization and salt-stratum balancing

The edge matrix was normalized at the individual LC-MS/MS run level using trimmed mean of M-values with singleton pairing (TMMwsp), following the edgeR TMMwsp formulation^45^. The reference run was selected as the run with the largest sum of square-root-transformed edge counts, and normalization factors were rescaled to have a geometric mean of 1. Runs with non-positive total assigned interlink mass were not eligible for this transformation.

For run *r*, let *L_r_* = ∑*_e_ C_er_* denote total assigned interlink mass, *f_r_* its TMMwsp normalization factor and *L*^∗^ = *L f* its effective library size. The scaled prior count was 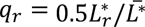 where *L̄*^∗^ is the mean effective library size across retained runs. Two run-level representations were calculated:

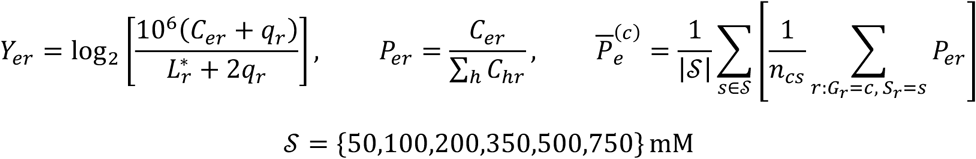

Here, *Y_er_* is the normalized log-counts-per-million value used for edge-level modeling, *P_er_* is the within-run composition of interlink evidence, *G_r_* and *S_r_* denote the condition and salt stratum of run *r*, respectively, and *n_cs_* is the number of retained runs in condition *c*and stratum *s*. TMMwsp factors were estimated once from the pooled edge-by-run matrix and were unchanged during label permutations because their calculation did not use condition labels.

The within-run compositions *P_er_* sum to 1 and were used for global compositional comparisons, condition-symmetric module construction and module evidence-share summaries. TMMwsp-normalized counts per million were not treated as a second independent measure of total PPI load because they are proportional to *P_er_*/*f_r_*. Accordingly, module-level values derived from these profiles were interpreted as relative interlink-evidence shares, not as absolute complex abundance, interaction stoichiometry or molecular occupancy.

Because the numbers of LC-MS/MS runs differed among salt strata, condition summaries were obtained by first averaging runs within each condition-by-stratum cell and then assigning equal weight to the six SCX strata, as shown in equation above. This prevented strata represented by more LC-MS/MS acquisitions from receiving greater influence in the condition comparison.

### Edge-level condition-association analysis

For each of the 213 protein-pair edges, normalized logCPM values were analyzed using a linear model containing condition and SCX salt stratum:

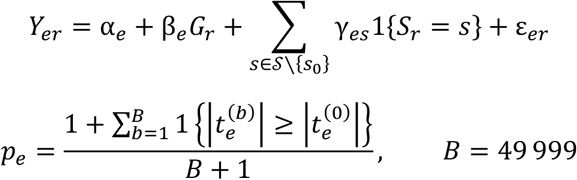

Here, *G_r_* = 1 for A636T and *G_r_* = 0 for WT, *s*_0_ is the reference salt stratum, *β_e_* is the estimated A636T-versus-WT log2 difference in normalized interlink evidence, 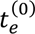 is the observed moderated *t*-statistic and 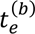 is the statistic under restricted permutation *b*.

Edge-specific residual variances were moderated across the complete 213-edge family using a scaled inverse-*χ*^2^ empirical-Bayes model analogous to the limma procedure. Empirical-Bayes prior hyperparameters were estimated from the observed pooled dataset and then held fixed for all permuted label assignments, so the same moderation reference was applied throughout the null calculation. Parametric moderated-model p-values were retained as diagnostics. The restricted-permutation values were used for primary edge-level inference.

Condition labels were permuted only among LC-MS/MS runs within the same SCX salt stratum, with the observed numbers of A636T and WT labels preserved within each stratum. This restriction retained the salt-fractionation structure but assumes that retained LC-MS/MS runs are technically exchangeable within a salt stratum. The resulting p-values were adjusted across all 213 tested edges using the Benjamini-Hochberg procedure. An absolute log _2_(1.5) = 0.585 effect threshold was used only as a descriptive effect annotation and was not used for prefiltering or significance testing. Because the scale of *β_e_* can depend on edge sparsity, this common threshold was not interpreted as an equivalent biological effect for all edges.

A pooled, condition-label-independent support weight was calculated for module construction and directional module summaries:

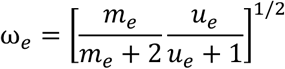

where *m_e_*is the total assigned CSM mass pooled across conditions and *u_e_* is the number of supporting LC-MS/MS runs. Mapping specificity was not included in the primary support weight because, under the unambiguous assignment rule in equation above, all assigned evidence has mapping specificity exactly equal to 1. The support weight did not enter edge-level p-value estimation.

### Global redistribution of protein-pair interlink evidence

Global redistribution of interlink evidence was evaluated using four complementary metrics. Jensen-Shannon divergence and weighted Jaccard distance were calculated from the equal- stratum condition profiles **P̄**^(A636T)^ and **P̄**^(WT)^. Weighted Jaccard distance was defined as one minus the sum of edgewise minima divided by the sum of edgewise maxima.

Graph-level differences were assessed independently using DeltaCon distance and network portrait divergence^53^. These analyses were restricted to the 197 loop-free heteromeric edges and used the same fixed 357-protein node universe in the observed and all permuted assignments. Positive edges were ranked separately in each condition by their equal-stratum mean compositions. To maintain identical graph density for every observed and permuted comparison, exactly *K* = 67 top-ranked positive edges were retained in each condition. This value was the largest common edge count feasible in both conditions across the complete restricted-permutation ensemble used for the forest-aware topology analysis. Deterministic, label-invariant ordering was used to resolve exact ranking ties. Sensitivity to boundary ties was evaluated using 200 randomized tie-resolution replicates.

Jensen-Shannon and weighted Jaccard statistics were evaluated using 49,999 restricted within-stratum permutations. DeltaCon and portrait divergence were evaluated using 4,999 restricted permutations because these metrics required repeated graph-distance calculations. Individual global p-values were upper-tail permutation values, and Holm correction was applied across the four prespecified global metrics.

A secondary four-metric omnibus statistic was calculated over the common set of graph-distance permutations. For each metric, the observed value and all permuted values were standardized using their pooled *B*_g_ + 1 mean and standard deviation, where *B*_g_ = 4,999. The omnibus statistic was the sum of the four squared standardized deviations. Its permutation p-value was calculated by comparing the observed omnibus with the omnibus values obtained from the same exchangeably standardized assignments. The omnibus was retained as a secondary global summary and did not replace interpretation of the four individual metrics.

### Contrast-blind consensus PPI-module construction

PPI modules were defined before differential module testing and without using observed edge-level fold changes, *t*-statistics or p-values. Only the 197 positive, loop-free heteromeric edges were eligible. In each of 200 bootstrap iterations, LC-MS/MS runs were sampled with replacement within each of the 12 condition-by-salt-stratum cells while preserving the original number of runs in each cell. Edge compositions were averaged within each resampled cell and then averaged equally across all 12 cells, producing a condition-symmetric pooled edge profile. Thus, condition membership was used only to prevent unequal cell sizes from biasing the pooled profile. The A636T-versus-WT contrast was not used to define modules.

Bootstrap edge weights were calculated as the pooled edge composition multiplied by 0.25 + 0.75*ω_e_*. A weighted partition was generated in each bootstrap using the Leiden RBConfiguration method with resolution parameter 1.0. For each pair of proteins, the proportion of bootstrap partitions in which the two proteins were assigned to the same community was recorded. A consensus graph was formed by connecting proteins with coassignment proportion of at least 0.50. Final modules were obtained by Louvain community detection on the weighted consensus graph; connected components were used as a fallback if the Louvain partition could not be obtained^54^. Module assignments were frozen before condition-association testing.

For module *M*, let ℰ*_M_* denote its internal heteromeric edges. Three complementary module statistics were calculated:

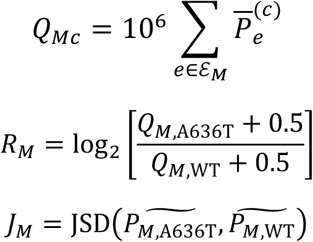

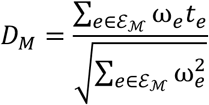

Here, *Q_Mc_* is the module’s relative interlink-evidence share expressed in parts per million, *R_M_* is its A636T-versus-WT evidence-share log ratio, **P**^]^*_Mc_* is the vector of internal edge shares renormalized to sum to 1 within module *M*, *J_M_* quantifies redistribution among a module’s internal edges, and *D_M_* summarizes the directionally concordant edge-level moderated statistics. The 0.5- p.p.m. pseudocount in *R_M_* was used solely to obtain a finite ratio when a module had no detected internal-edge evidence in one condition. Neither *Q_Mc_* nor *R_M_* was interpreted as absolute complex abundance.

Module inference used 49,999 restricted within-stratum permutations with the observed consensus-module assignments held fixed. The evidence-share ratio and directional statistic were evaluated two-sided, whereas JSD was evaluated in the upper tail. For assignment *b* = 0, …, *B*, where *b* = 0 denotes the observed assignment, the transformed statistics were:

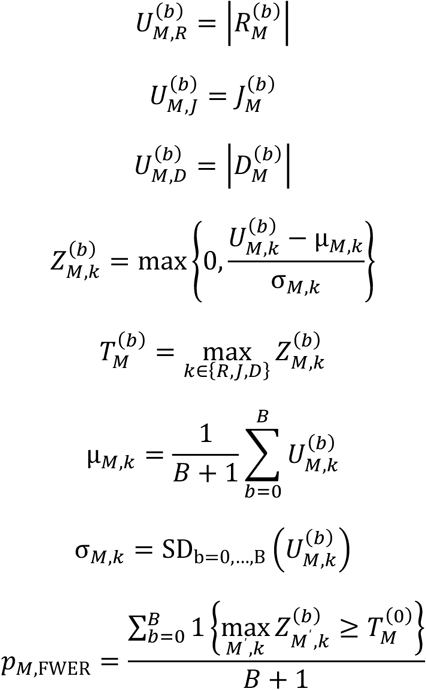

For every module–metric combination, *μ_M_*_,*k*_ and *σ_M_*_,*k*_ were calculated exchangeably from all *B* + 1 values, including the observed assignment. Module-metric combinations with zero or undefined variance were omitted from the omnibus rather than assigned a numerical *Z*-score. This is relevant, for example, when a module contains only one internal edge and its internal- composition JSD is identically zero.

Joint familywise error was controlled by comparing 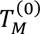 with the permutation distribution of the maximum 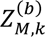 across every tested module and all three statistic families. The resulting joint maxT value was the primary module-level adjusted p-value. Metric-specific maxT values were retained as secondary outputs. Module direction and technical stability were additionally evaluated after omitting each SCX salt stratum in turn; these leave-one-stratum-out analyses recomputed module effect summaries while retaining the previously defined module membership.

### Connected-subnetwork analysis

A network-based statistic was used to identify connected sets of heteromeric edges showing concordant condition-associated remodeling. For each direction *d*, where *d* = 1 denoted A636T- higher evidence and *d* = −1 denoted WT-higher evidence, edges satisfying *dt_e_* > *τ* were selected at moderated-*t* thresholds *τ* ∈ 1.5,2.0,2.5,3.0. Connected components were identified among the selected edges, and component mass was defined as:

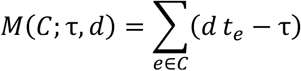

Components producing identical edge sets at multiple thresholds were deduplicated, retaining the representation with the largest component mass. For each of 49,999 restricted permutations, the maximum component mass across both directions and all thresholds was recorded. Component-level familywise-error-rate p-values were obtained from this maximum-null distribution. Network-based-statistic significance was interpreted at the connected-component level and did not imply that each constituent edge was individually significant.

### Forest-aware, density-matched network-topology analysis

A separate topology analysis evaluated whether the organization of the interlink-evidence network differed between conditions across matched graph densities. The analysis used the fixed union of 357 proteins and the 197 loop-free heteromeric edges. Proteins not incident to a retained edge at a particular density remained in the graph as isolates. Edges were ranked separately in A636T and WT by equal-stratum mean composition, and exactly the same number of top-ranked positive edges was retained in the two conditions.

Descriptive topology curves were calculated for every retained-edge count from *k* = 0 to the observed common ceiling of 103 edges. Formal permutation inference was restricted to 31 prespecified points spanning *k* = 0 to the permutation-feasible ceiling *K* = 67. Results beyond *k* = 67 were treated as descriptive only. Sensitivity to ties crossing an edge-retention boundary was assessed using 200 randomized tie-resolution replicates. The run manifest confirmed 172 runs, six permutation strata, 357 fixed nodes, 197 eligible edges and 49,999 restricted permutations.

The prespecified inferential family comprised ten topology metrics: active-protein fraction; largest-component fraction relative to the fixed node universe and to active proteins; binary global efficiency relative to the fixed universe and to active proteins; fixed-universe degree centralization; evidence-strength Gini coefficient relative to the fixed universe and to active proteins; normalized entropy of active-component sizes; and active-component susceptibility after exclusion of the largest component. Additional component-size, isolate, path-length, diameter, degree, evidence-strength, leaf, branch-node and cycle-rank measures were reported descriptively.

For metric *m* and assignment *b*, the condition-difference curve and whole-curve statistic were:

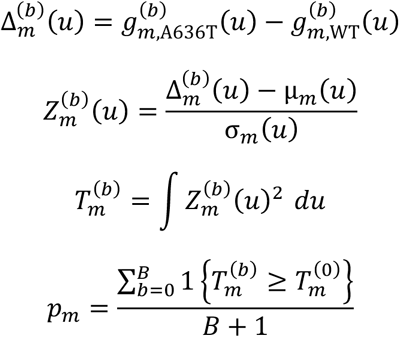

where *u* = *k*/*K*. The reference mean *μ_m_*(*u*) and standard deviation *σ_m_*(*u*) were calculated from all *B* + 1 assignments, including the observed assignment. Thus, observed and permuted curves were standardized by the same exchangeable rule. Only filtration positions that were finite for every assignment and had nonzero pooled variance contributed to the test. Holm correction was applied across the ten prespecified topology metrics.

All observed filtration graphs were forests over the evaluated density range. Triangle- dependent clustering coefficients and transitivity were therefore marked as not estimable for the observed graphs rather than assigned zero, and they were excluded from the primary inferential family. Separately optimized modularity was also excluded from primary topology inference because independently optimized A636T and WT partitions do not test correspondence of module assignments.

### Analysis of condition-restricted interlink detection

Binary detection was analyzed separately from quantitative interlink evidence because sparse sampling can produce edges detected in only one condition. For edge *e*and run *r*, occupancy was defined as *O_er_* = **1***C_er_* > 0. Occupancy was averaged first within each condition-by-stratum cell and then equally across the six salt strata. Condition differences were evaluated using 49,999 restricted within-stratum permutations, and permutation p-values were adjusted across the complete 213-edge family using the Benjamini-Hochberg procedure.

Connected patterns of differential occupancy were evaluated using a presence-based network statistic. For each edge, occupancy differences were converted to permutation- standardized *Z*-scores using the pooled observed-plus-permutation reference. The thresholds 1.5, 2.0, 2.5 and 3.0 therefore applied to the absolute standardized occupancy difference, not to raw occupancy differences. Component mass and familywise-error control were calculated across both directions and all four thresholds using the same maximum-component principle as the quantitative network-based-statistic analysis.

Technical robustness of condition-restricted detections was evaluated using 5,000 salt- stratum-balanced depth-matching and rarefaction iterations, leave-one-run-out analyses and leave-one-stratum-out analyses. These procedures reassessed the detection class and direction under reduced or balanced technical sampling; they did not convert SourceStem runs into biological replicates. Edges detected only in one specimen were described as “detected only in A636T” or “detected only in WT,” rather than as gained or lost interactions, because non-detection does not establish biological absence.

### Protein-abundance adjustment and prior-evidence annotation

Protein-abundance adjustment was performed as a secondary sensitivity analysis and did not enter the primary edge, module, connected-subnetwork or topology tests. For edges whose two endpoints were quantified in the independent TMT proteomic dataset, the BLIS edge log2 difference *β_e_* was modeled using Huber robust regression as a function of the summed TMT log2 abundance changes of the two endpoint proteins and their pair-level mean abundance. The residual from this model was termed the abundance-adjusted XL residual. Positive residuals indicated more A636T-associated interlink evidence than predicted from the endpoint abundance covariates, whereas negative residuals indicated less. Because the TMT and BLIS measurements were obtained from different specimen sets and the TMT covariates were themselves measured with error, these residuals were used for descriptive prioritization and were not described as abundance-independent effects.

Canonical protein pairs were compared with BioGRID records to annotate previously reported physical interactions. BioGRID overlap was treated as prior-evidence support rather than experimental validation because database coverage is uneven and favors extensively studied proteins. Structural compatibility and crosslink-supported modeling were likewise reported as orthogonal interpretive annotations and did not modify permutation p-values.

### Technical-evidence prioritization

Inferential quantities were retained at their appropriate analytical level and were not combined by taking the minimum across unrelated testing families. For a dyadic PPI, the primary adjusted value was the edge-level permutation Benjamini-Hochberg *q* value. For a predefined consensus module, the primary adjusted value was the joint module maxT familywise-error-rate value. For a network-based-statistic component, the component-level familywise-error-rate value was reported. Membership of an edge in a significant component was recorded as component-level support and was not substituted for the edge’s own adjusted p-value.

Recurrence across CSMs, LC-MS/MS runs and SCX strata; leave-one-run-out and leave-one-stratum-out stability; rarefaction stability; resolved residue-pair evidence; protein-abundance adjustment; BioGRID overlap; and structural compatibility were reported as separate evidence dimensions. Any resulting evidence tiers were interpreted as technical-prioritization categories, not as additional levels of statistical significance. Biological or disease relevance was considered only after technical ranking.

### Software and reproducibility

All BLIS 1.0 analyses were implemented in Python and executed under Ubuntu 22.04 in Windows Subsystem for Linux 2 using Python v.3.12.13. The principal computational environment contained NumPy v.2.4.6, pandas v.3.0.3, SciPy v.1.17.1, Matplotlib v.3.11.0, Biopython v.1.87, NetworkX v.3.6.1, igraph v.1.0.0, leidenalg v.0.12.0, openpyxl v.3.1.5 and gprofiler-official v.1.0.0. Large language models (LLMs) were used as assistive tools during BLIS development for refinement of mathematical formulations, auditing and debugging of analysis scripts, and code annotation and documentation. All LLM-assisted outputs were manually reviewed, tested and, where applicable, independently verified before incorporation into the final analysis workflow.

### Structural and network analysis software

STRING was used for protein-protein interaction network and functional-enrichment analyses. Protein and protein-complex structures were predicted using the AlphaFold 3 web server. Crosslink-guided structural modeling was performed with AlphaLink2 on Northwestern University’s QUEST high-performance computing cluster using experimentally identified interprotein crosslinks as spatial restraints. Candidate models were assessed using model-confidence metrics and agreement with experimental crosslink restraints. Structural models were inspected and rendered in UCSF ChimeraX v1.10, including visualization of protein chains and crosslinked residues, inter-residue distance measurements and preparation of structural figures.

## Data availability

The MS data generated in this study will be deposited in a publicly accessible repository upon acceptance of the manuscript.

## Code availability

The BLIS 1.0 source code and analysis scripts will be available on GitHub upon acceptance of the manuscript.

## Ethics declarations

The authors declare no competing financial interests.

## Supporting information

BLIS_Additional information

## Acknowledgements

This work was supported by R01 MH130428 to AC and JNS, R01 MH099114 to AC, R01 NS115471 to AC, R01 EY032506 to AC, R01 AG078796 and S10 OD032464 to JNS. This research was supported in part through the computational resources and staff contributions provided for the Quest high performance computing facility at Northwestern University which is jointly supported by the Office of the Provost, the Office for Research, and Northwestern University Information Technology.

## Contributions

Y-Z W conceived the study, designed the experiments, performed the experiments, developed the computational code, analyzed the data, and wrote the original draft of the manuscript. JX bred the animals and performed brain fixation and sectioning. JX, JNS, and AC contributed to the interpretation of the results and critically reviewed and edited the manuscript. All authors reviewed and approved the final manuscript.

---

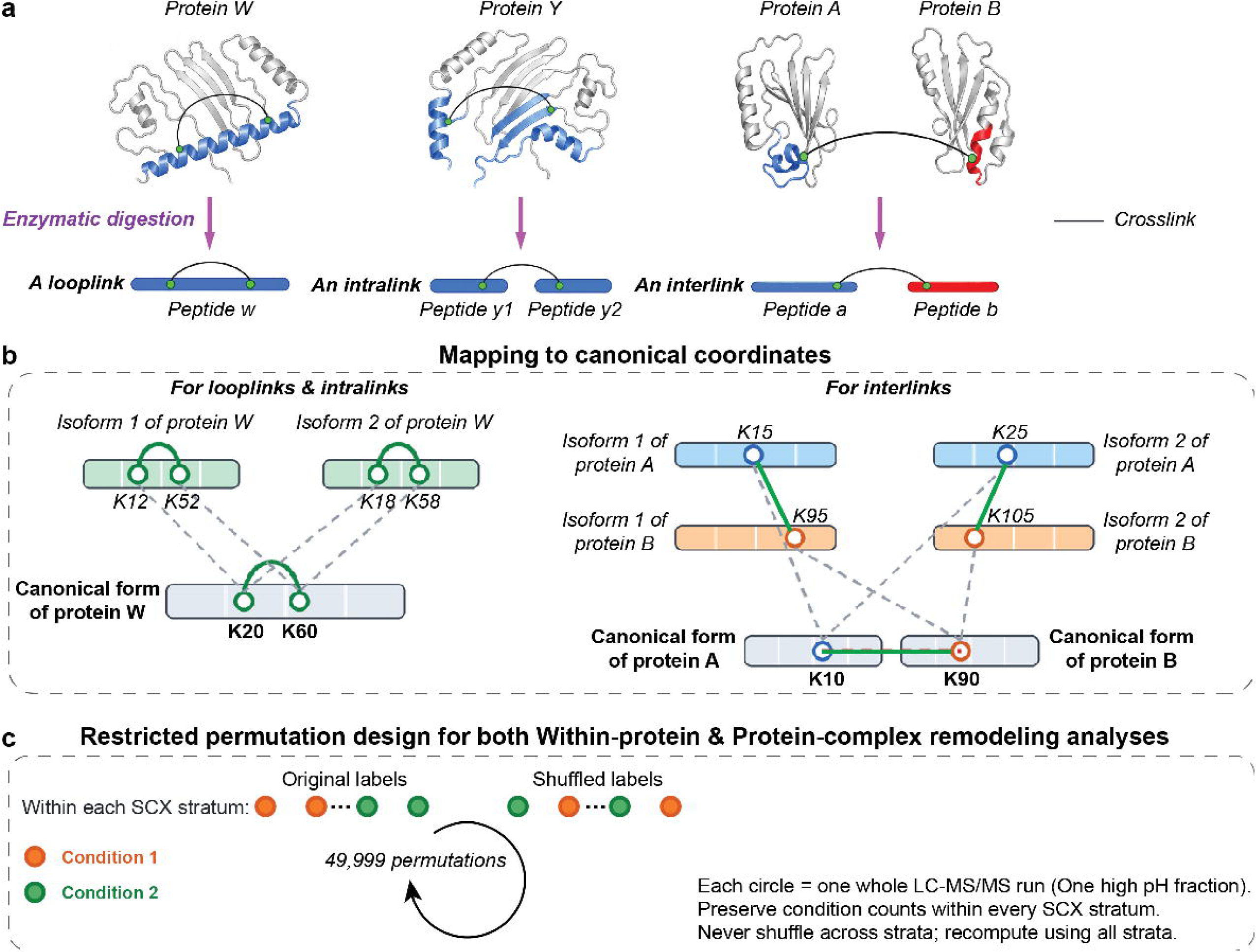

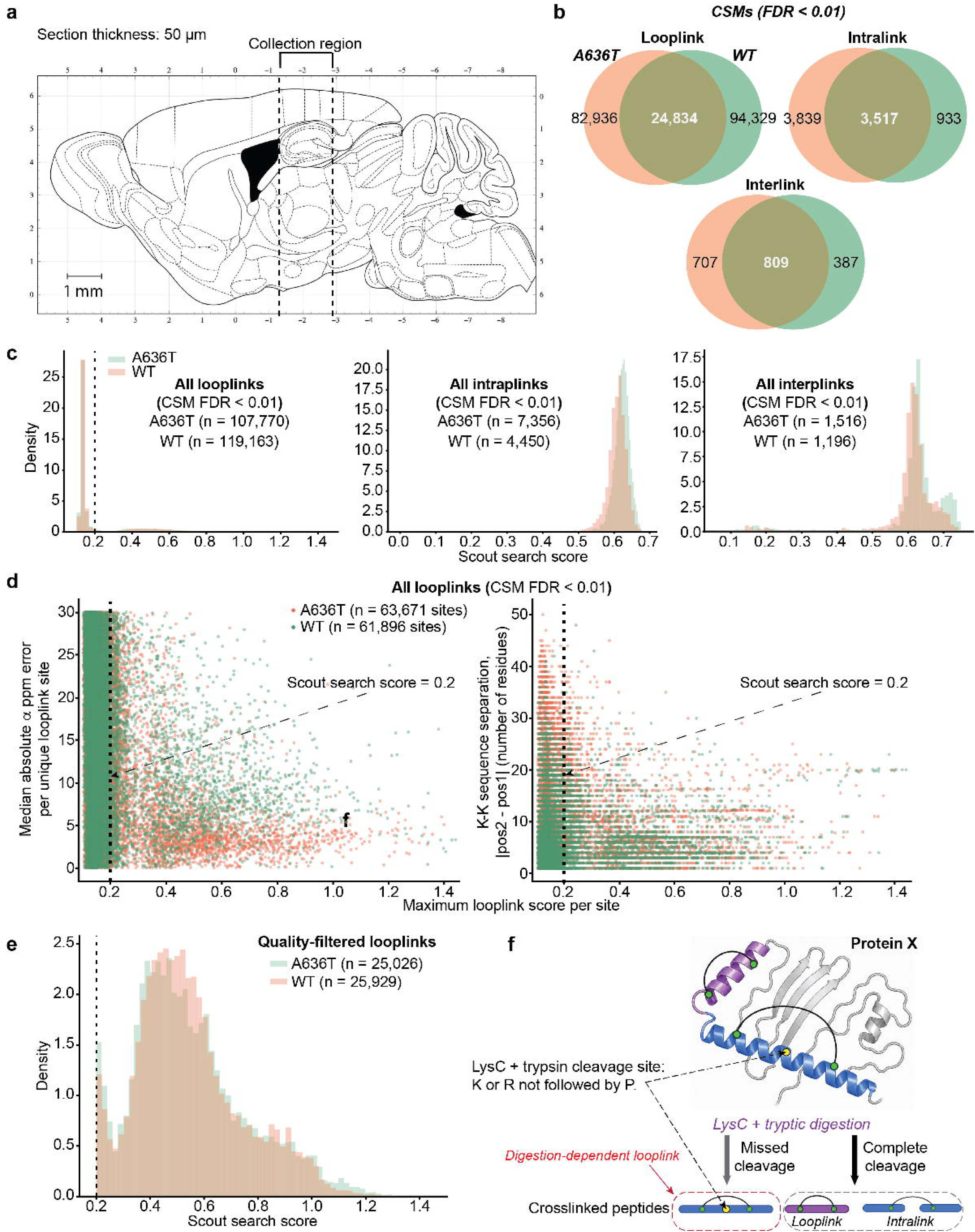

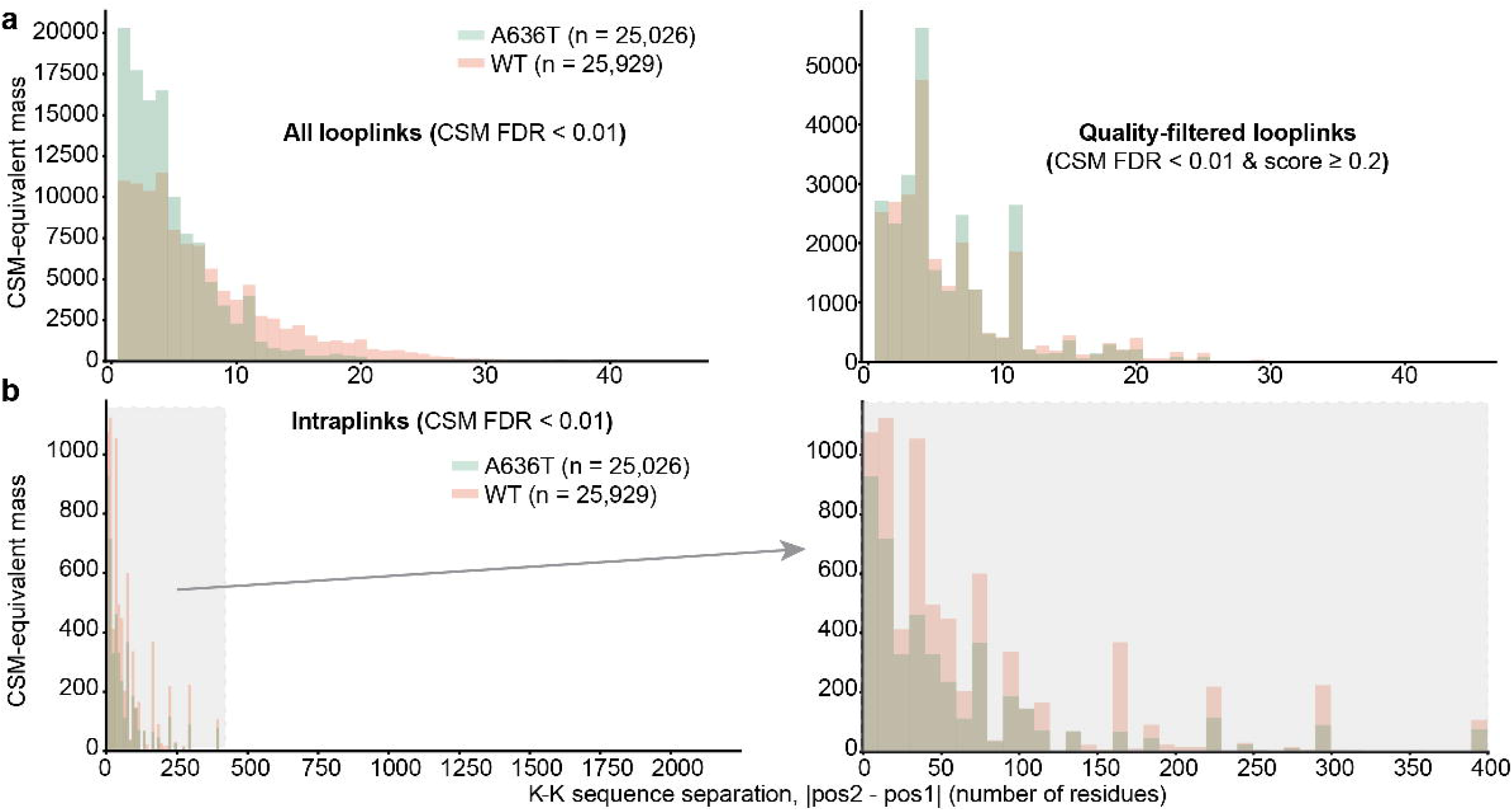

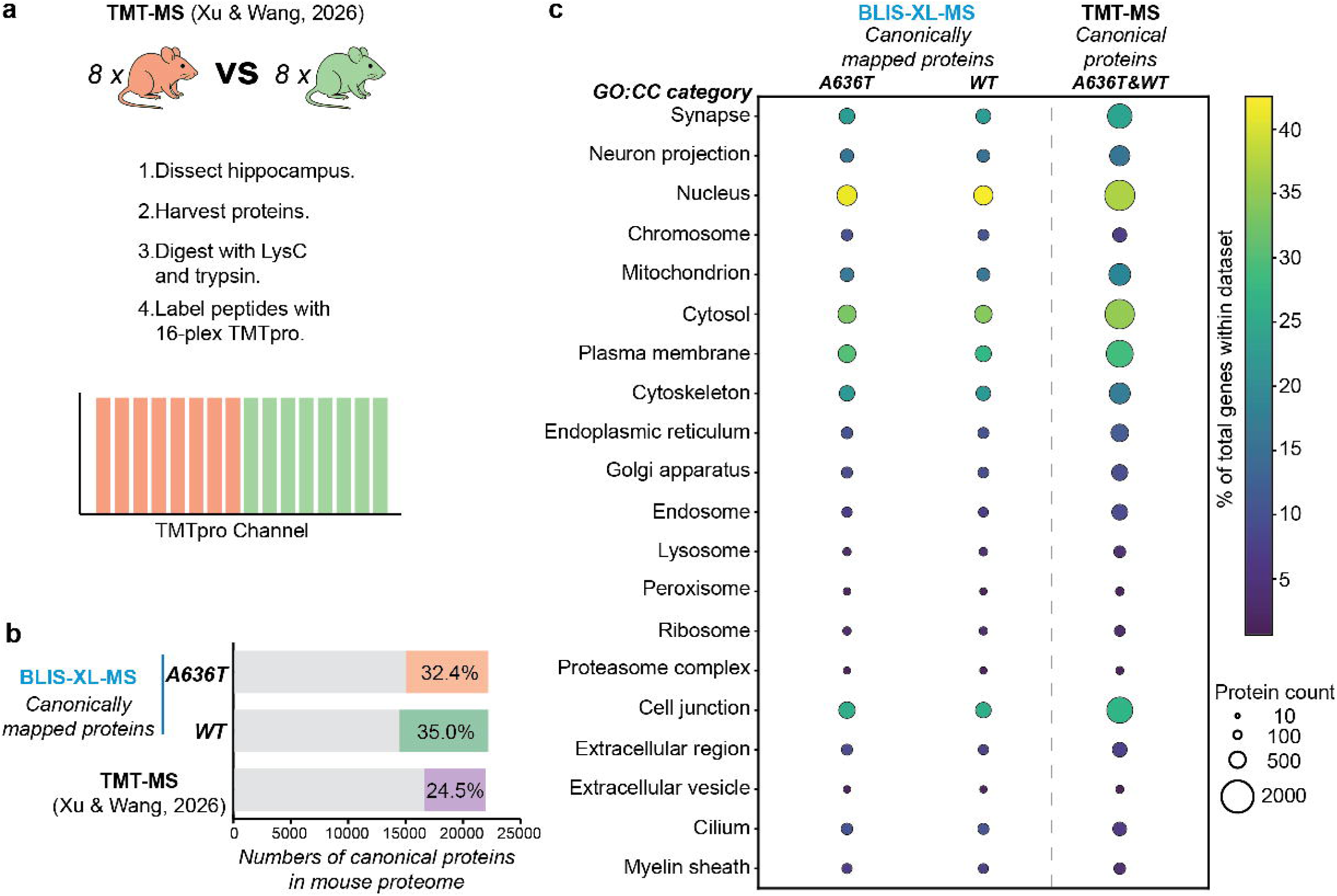

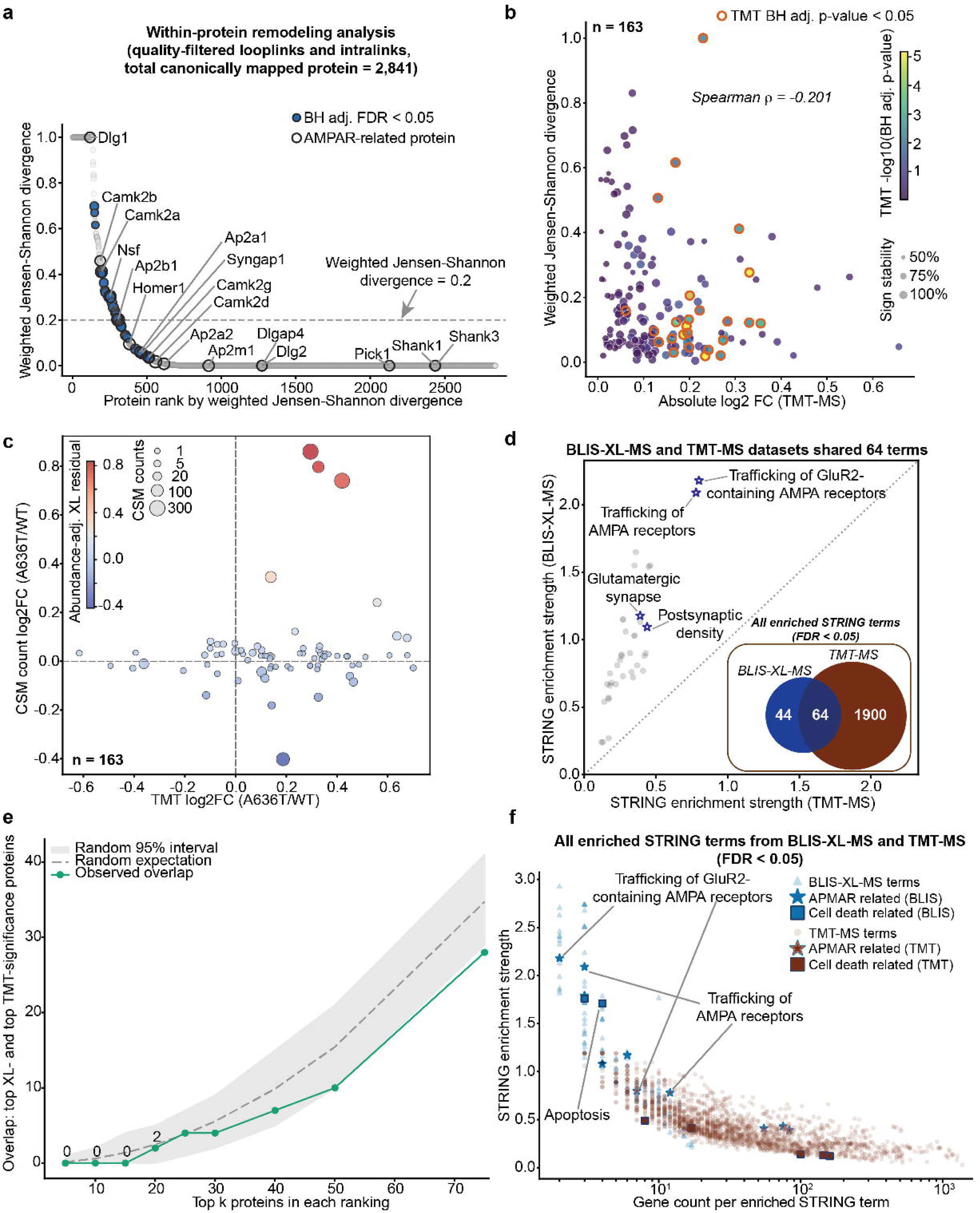

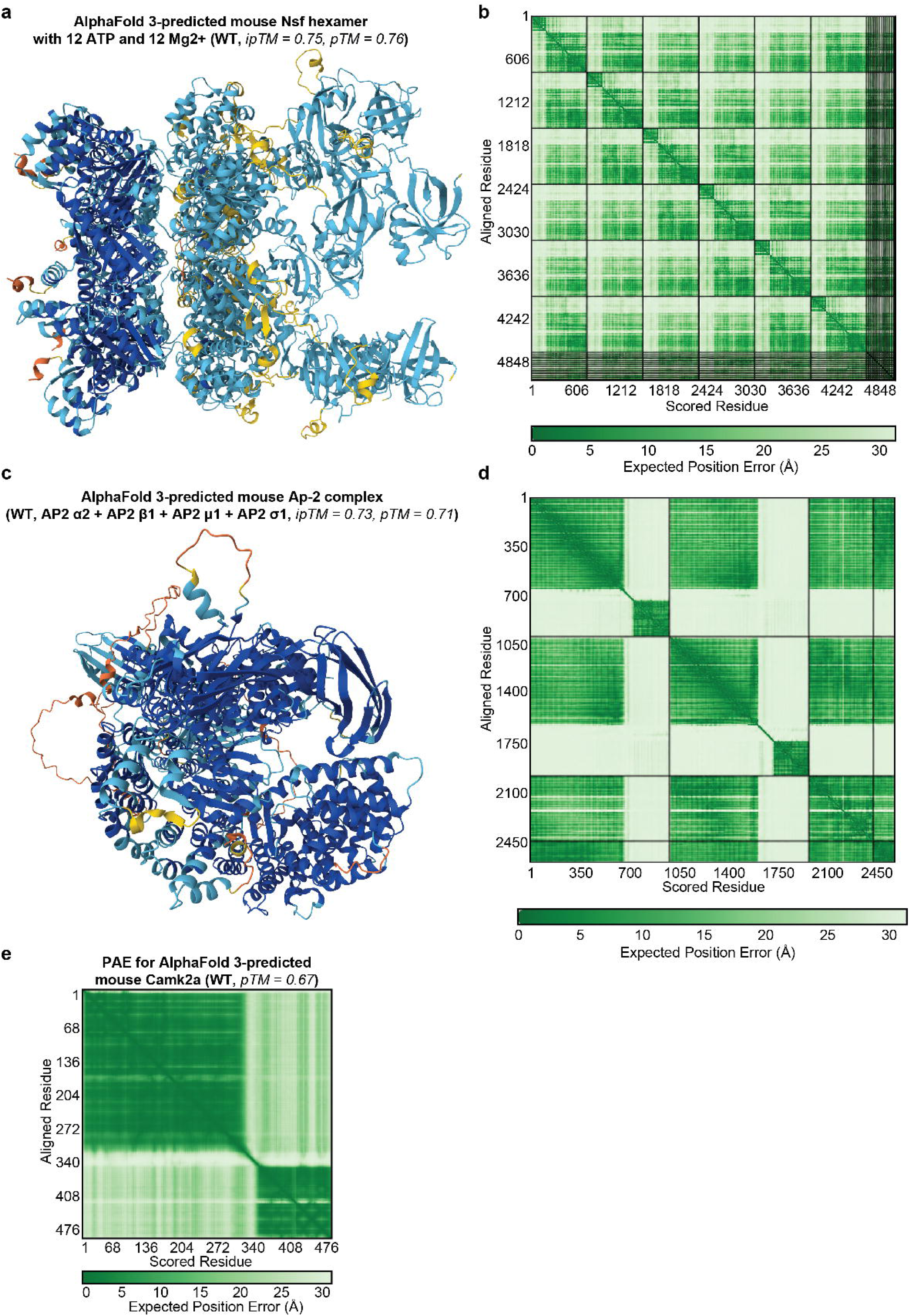

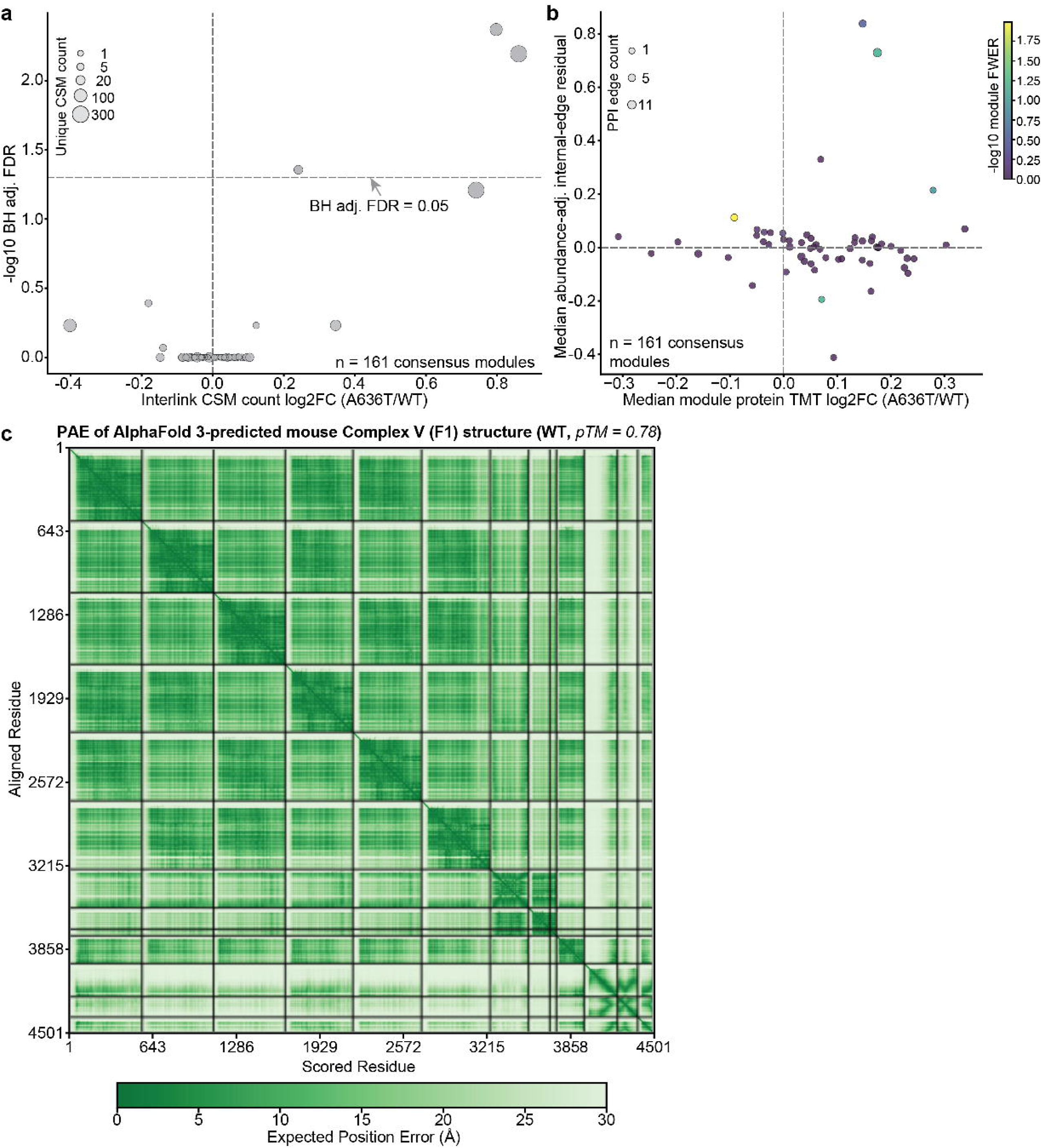

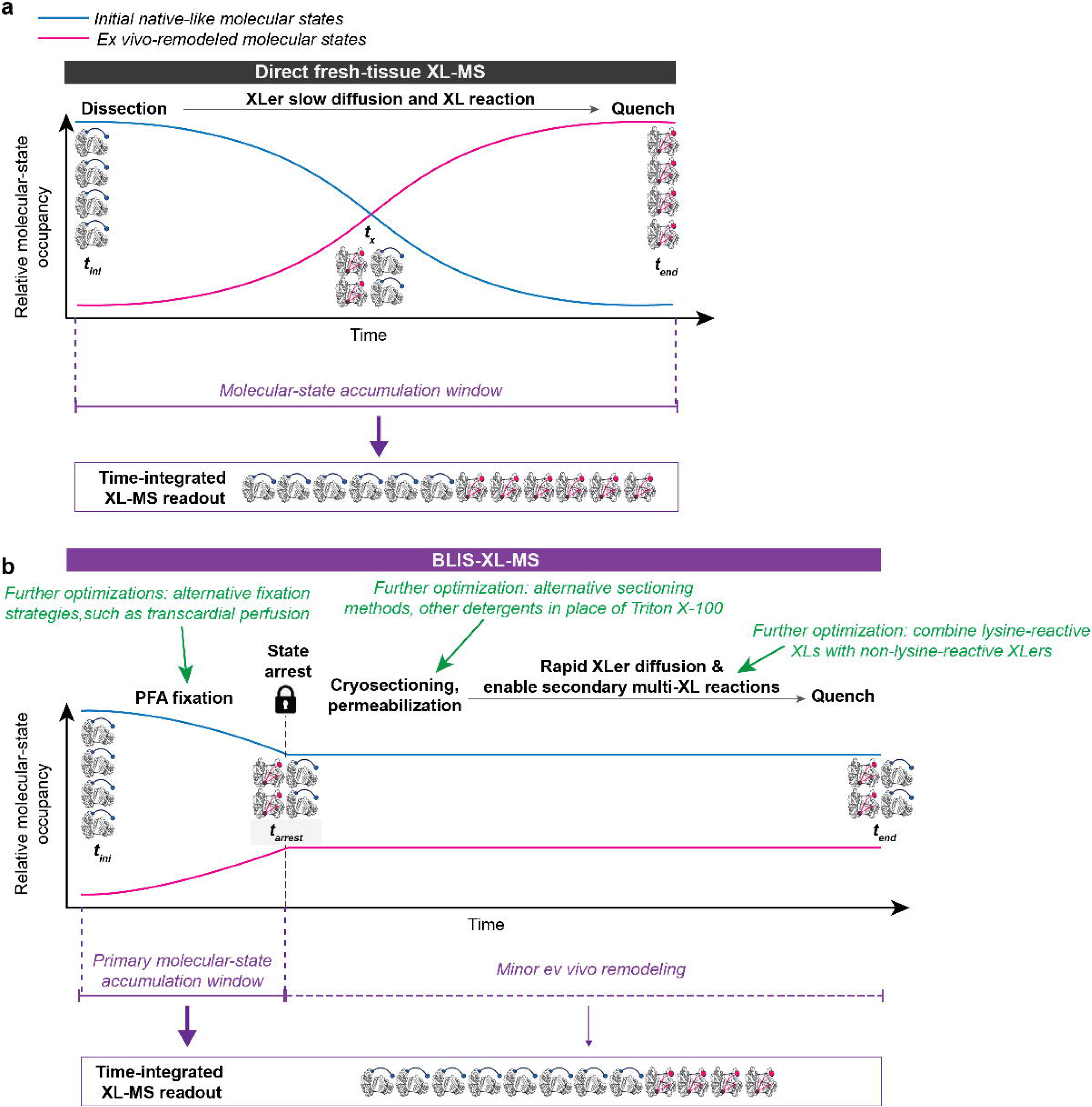

