## Supplementary material for "Brain Landscape In Situ Crosslinking Mass Spectrometry (BLIS-XL-MS) Enables Global Analysis of Protein Structural Remodeling in the Brain": BLIS_Additional information

**Extended data**

**Extended Data Fig.1 Crosslink classes, canonical-coordinate mapping and restricted permutation strategy used in BLIS 1.0.**

**a**, Schematic representation of the three crosslink classes used in BLIS analyses. Looplinks connect two lysine residues within the same peptide, intralinks connect two peptides derived from the same protein, and interlinks connect peptides derived from different proteins.

**b**, Mapping of crosslinked lysine residues from protein isoforms to canonical protein coordinates. For looplinks and intralinks, isoform-specific lysine positions are remapped to a common canonical sequence to define comparable within-protein residue pairs. For interlinks, lysines from each interacting protein are independently mapped to their corresponding canonical sequences, yielding canonical protein-pair and residue-pair coordinates.

**c**, Restricted permutation design used for both within-protein and protein-complex remodeling analyses. Condition labels are shuffled only among complete LC-MS/MS runs within the same SCX salt stratum, while preserving the original number of runs assigned to each condition within every stratum. Labels are never exchanged across SCX strata, and the full analysis statistic is recomputed after each shuffle using evidence from all strata. Empirical p-values are estimated from 49,999 restricted permutations.

Relates to **Fig.1b.**

**Extended Data Fig.2 Quality assessment and filtering of looplinks for within-protein remodeling analysis.**

**a,** The brain section region spans the indicated brain region which cover hippocampus. Relates to **Fig.2a.**

**b,** Overlap of Scout-accepted crosslink evidence between A636T and WT. Venn diagrams show matched looplink, intralink and interlink identities at CSM FDR < 0.01. Looplinks comprised 82,936 A636T-specific, 24,834 shared and 94,329 WT-specific CSM assignments; intralinks comprised 3,839 A636T-specific, 3,517 shared and 933 WT-specific assignments; and interlinks comprised 707 A636T-specific, 809 shared and 387 WT-specific assignments. The substantially different overlap patterns among link classes motivated link-class-specific quality assessment and downstream treatment.

**c,** Distributions of Scout search scores for looplink, intralink and interlink CSMs accepted at a CSM-level FDR < 0.01. Looplinks showed a distinct score distribution, with a large population concentrated below a Scout score of 0.2, whereas intralinks and interlinks were predominantly distributed at higher scores. The vertical dashed line indicates a Scout score of 0.2.

**d,** Site-level quality characteristics of all looplinks passing CSM FDR < 0.01. Each point represents a unique looplink CSMs, summarized across supporting CSMs (A636T, *n* = 63,671 CSMs; WT, *n* = 61,896 CSMs). *Left*, maximum Scout search score for each site plotted against its median absolute α-peptide precursor-mass error. Low-scoring looplink sites showed a substantially broader distribution of absolute mass errors, including enrichment for large ppm errors, consistent with reduced confidence in peptide assignment. *Right*, maximum Scout search score plotted against K-K sequence separation, defined as |pos2 − pos1|. Low-scoring evidence was also strongly concentrated at short sequence separations, although short-range links were not restricted to the low-score population. The vertical dashed line denotes the looplink quality-filtering threshold at a Scout score of 0.2.

**e,** Scout score distributions after removal of low-scoring looplinks. Quality-filtered looplinks retained a substantial population spanning approximately 0.2-1.2 in Scout score and showed closely overlapping distributions between A636T (*n* = 25,026 CSMs) and WT (*n* = 25,929 CSMs). Thus, filtering removed the dominant low-score population while retaining a broad range of higher-confidence looplink evidence.

**f,** Schematic illustrating the dependence of looplink versus intralink classification on local proteolytic cleavage. For two crosslinked residues within the same protein, complete cleavage at an intervening LysC/trypsin-compatible site can generate two crosslinked peptides and therefore an intralink. A missed cleavage at the same intervening site retains both linked residues within a single peptide, causing the corresponding restraint to be observed as a looplink. Such missed-cleavage-derived looplinks therefore represent digestion-dependent peptide products rather than a fundamentally different class of within-protein structural restraint.

**b-f** relate to **Fig.2d.**

**Extended Data Fig. 3 Sequence-separation distributions of looplinks and intralinks in A636T and WT brains.**

**a**, Distribution of lysine-lysine (K-K) sequence separation for looplinks in A636T and WT samples, weighted by CSM-equivalent mass. *Left*, all looplinks passing CSM FDR < 0.01; *right*, quality-filtered looplinks additionally requiring a Scout score ≥ 0.2.

**b**, Distribution of K-K sequence separation for intralinks passing CSM FDR < 0.01. *Left*, full sequence-separation range; right, enlarged view of the shaded 0–400-residue region. Histograms show A636T (green; n = 25,026) and WT (orange; n = 25,929) evidence distributions. Sequence separation was calculated as |pos2 − pos1| after mapping crosslinked lysines to protein sequence coordinates.

Relate to **Fig.2d.**

**Extended Data Fig. 4 Proteome coverage and subcellular representation of BLIS-XL-MS compared with quantitative TMT-MS.**

**a,** Experimental design of the reference quantitative proteomics dataset (Xu & Wang, 2026). Hippocampi from eight heterozygous GluA1^A636T^ mice and eight WT littermates were dissected, proteins were extracted and sequentially digested with LysC and trypsin, and peptides were labeled with 16-plex TMTpro for quantitative LC-MS/MS analysis.

**b,** Proteome coverage of BLIS-XL-MS and TMT-MS relative to the canonical mouse proteome. Canonically mapped BLIS-XL-MS proteins represented 32.4% and 35.0% of the mouse proteome in A636T and WT samples, respectively, compared with 24.5% for the TMT-MS dataset.

**c**, Gene Ontology Cellular Component (GO:CC) representation of canonically mapped proteins detected by BLIS-XL-MS in A636T and WT samples and by TMT-MS. Circle size indicates the number of proteins assigned to each GO:CC category, and color indicates the percentage of proteins within each dataset assigned to that category. BLIS columns represent genes mapped from all quality-filtered XLs in A636T and WT separately. The TMT-MS column represents all genes quantified across the A636T and WT TMT samples, rather than only differentially abundant proteins. The similar representation of major compartments, including synapse, neuronal projection, mitochondrion, cytosol, plasma membrane and intracellular organelles, indicates broad subcellular sampling by BLIS-XL-MS without an obvious compartment-level coverage bias relative to conventional proteomics.

Relate to **Fig.3a-b.**

**Extended Data Fig. 5 Benchmarking BLIS-XL-MS within-protein remodeling against TMT-MS abundance profiling.**

**a,** Proteins detected by the BLIS-XL-MS within-protein analysis using quality-filtered looplinks and intralinks were ranked by weighted JSD (0-1) between A636T and WT (*n* = 2,841 canonically mapped proteins). Each point represents one canonically mapped protein. Proteins significant after BH correction (BH-adjusted FDR < 0.01) are shown in blue, and AMPAR-related proteins are indicated by open circles. Selected AMPAR-associated and synaptic proteins are labelled. The horizontal dashed line indicates weighted JSD = 0.2. Several synaptic and AMPAR-associated proteins, including Dlg1, Camk2a, Camk2b, Nsf, Ap2b1, Homer1, and Syngap1, were among the highest-ranked remodeling candidates, indicating that the BLIS within-protein remodeling analysis captured biologically coherent structural changes rather than a random set of proteins.

**b,** Comparison of BLIS-JSD with protein-abundance changes measured by the independent TMT-MS dataset. Weighted JSD is plotted against the absolute TMT log2 fold change (A636T/WT) for the 163 BLIS-tested proteins. Point color represents $-\log_{10}$of the BH-adjusted TMT p-value, and orange outlines identify proteins significant by TMT-MS (BH-adjusted p-value < 0.05). Point size denotes the resampling-based stability of the direction of the TMT abundance effect, with larger points indicating greater sign stability. Weighted JSD showed only a weak inverse relationship with the magnitude of protein-abundance change (Spearman’s $\rho=-0.201$), indicating that the strongest structural-profile differences were not preferentially associated with the largest abundance changes. The TMT-MS measurements were used as an orthogonal abundance reference rather than as a direct replicate-level validation of the BLIS measurements.

**c,** Relationship between protein-abundance changes measured by TMT-MS and changes in within-protein CSM abundance for proteins eligible for BLIS statistical testing (n = 163). The x axis shows TMT log2 fold change (A636T/WT), and the y axis shows CSM-count log2 fold change (A636T/WT). Point size represents the number of supporting CSMs. Point color represents the abundance-adjusted XL residual, defined as the deviation of the observed CSM-count change from that expected after accounting for protein abundance; positive (red) values indicate greater XL evidence than expected from abundance, whereas negative (blue) values indicate less. Dashed lines indicate no change. CSM-count changes showed limited overall concordance with protein-abundance changes, although pronounced abundance differences could influence crosslink counts for individual proteins.

**d,** Comparison of STRING enrichment strength between the BLIS within-protein remodeling analysis and TMT-MS abundance analysis for terms significantly enriched in both datasets. Each point represents one of the 64 shared enriched terms; the x and y axes show STRING enrichment strength for TMT-MS and BLIS-XL-MS, respectively. The dotted diagonal indicates equal enrichment strength in the two analyses. Blue stars denote selected AMPAR- and synapse-associated terms, *Inset*, overlap of all significantly enriched STRING terms (FDR < 0.05): 108 terms were detected from the BLIS conformational-remodeling candidates, of which 64 were shared with TMT-MS and 44 were BLIS-specific; TMT-MS yielded 1,964 enriched terms, including 1,900 not present in the BLIS enrichment set.

**e,** Rank-overlap analysis between proteins prioritized by BLIS within-protein remodeling and by TMT-MS abundance change. The green line shows the observed number of proteins shared between the top *k* proteins in the two rankings. The grey dashed line and shaded region indicate the random expectation and its 95% interval, respectively. Numbers indicate the observed overlap at selected values of *k*. The limited overlap at stringent ranking thresholds indicates that proteins prioritized by structural remodeling are largely distinct from those prioritized by abundance change.

**f,** Comparison of all significantly enriched STRING terms identified from BLIS-XL-MS and TMT-MS (FDR < 0.05). Each symbol represents one enriched term, plotted according to the number of genes assigned to the term and STRING enrichment strength. Triangles denote BLIS-XL-MS terms and circles denote TMT-MS terms. AMPAR-related terms are highlighted by stars and cell-death-related terms by squares. Representative AMPAR-trafficking and apoptosis terms are labelled. BLIS-XL-MS yielded a smaller, more focused enrichment set than TMT-MS, with AMPAR-trafficking terms among the prominent BLIS enrichments.

Relate to **Fig.3b.**

**Extended Data Fig.6 AlphaFold 3 model-confidence assessment of three AMPAR-trafficking proteins.**

**a**, AlphaFold 3-predicted structure of the mouse NSF hexamer in an ATP-bound state, modeled with 12 ATP molecules and 12 Mg2+ ions. The selected WT model had an interface predicted TM score (ipTM) of 0.75 and a predicted TM score (pTM) of 0.76. The hexameric model is colored by residue-level AlphaFold 3 confidence (pLDDT), with blue representing higher-confidence regions and yellow-orange representing lower-confidence regions. The nucleotide-bound model was used to provide structural context for interpreting BLIS-derived remodeling evidence in NSF.

**b**, Predicted aligned error (PAE) matrix for the NSF model shown in **a**. Rows and columns correspond to aligned and scored residues, respectively, across the six NSF protomers. Darker green indicates lower predicted positional error and therefore greater confidence in the relative placement of the corresponding residues, whereas progressively lighter colors indicate greater uncertainty. The repeated diagonal blocks reflect high confidence within individual NSF protomers, while the extensive low-PAE signal between protomers supports the overall relative organization of the modeled hexamer. Black lines delineate protomer boundaries.

**c**, AlphaFold 3-predicted structure of the mouse AP-2 adaptor complex comprising AP2 α2, AP2 β1, AP2 μ1 and AP2 σ1 subunits. The selected WT model had an ipTM of 0.73 and a pTM of 0.71. The model is colored by residue-level confidence as in a. This complex includes AP2B1, one of the proteins prioritized by the BLIS within-protein remodeling analysis, and was modeled to place the observed remodeling in the context of the assembled AP-2 complex rather than an isolated subunit.

**d**, PAE matrix for the AP-2 complex shown in **c**. The four major diagonal regions correspond to the four modeled AP-2 subunits. Low PAE within these regions indicates high confidence in the local folds of the individual subunits, whereas variation in the off-diagonal blocks reflects differing confidence in their relative positioning and intersubunit organization. The matrix therefore distinguishes well-constrained portions of the AP-2 core from regions for which the relative domain or subunit arrangement is less certain. Black lines indicate subunit boundaries.

**e**, PAE matrix for an AlphaFold 3-predicted mouse CaMKIIα (CAMK2A) monomer (WT; pTM = 0.67). The matrix shows high internal confidence within the major structured regions, including the N-terminal kinase-containing region and the C-terminal association/hub region, but increased uncertainty in their relative positioning across the intervening regulatory/linker region. This pattern is consistent with a model in which the individual structured domains are more confidently predicted than their interdomain orientation. Unlike NSF and AP-2, the full CaMKIIα oligomeric complex was not modeled because the multimeric assembly exceeded the 5,000-token input limit of the AlphaFold 3 implementation used here. We therefore modeled a single CaMKIIα protomer and used its predicted structure and PAE profile to assess confidence in the major intra-subunit domains and their relative organization, rather than to infer the architecture of the complete CaMKIIα holoenzyme.

PAE is shown in ångströms for **b**, **d** and **e**, with darker green indicating lower expected positional error. Relate to **Fig.3c.**

**Extended Data Fig. 7 AlphaFold 3 assessment of mitochondrial Complex V.**

**a**, Differential interlink evidence across the 161 condition-blind consensus PPI modules defined from the BLIS interlink network. Each point represents one consensus module. The x axis shows the module-level interlink CSM-count log2 fold change (A636T/WT), and the y axis shows -log10 of the BH-FDR. Point size represents the number of unique CSMs contributing to the module. The vertical dashed line denotes no change between genotypes, and the horizontal dashed line denotes BH-FDR = 0.05. Several modules showed increased interlink evidence in A636T, including the ATP synthase-centered modules identified among the strongest remodeling signals.

**b**, Relationship between protein-abundance change and abundance-adjusted PPI remodeling across the same 161 consensus modules. The x axis shows the median TMT-MS protein log2 fold change (A636T/WT) for proteins within each module, and the y axis shows the median abundance-adjusted internal-edge residual, quantifying PPI remodeling remaining after accounting for the abundance changes of the proteins contributing to each interaction. Point size indicates the number of internal PPI edges in the module, and point color represents -log10 of the module-level family-wise error rate (FWER) obtained from the permutation-based module analysis. Dashed lines indicate zero abundance change and zero abundance-adjusted PPI residual. Most modules clustered near zero residual despite variation in protein abundance, whereas a small subset showed pronounced positive or negative residuals, indicating interaction remodeling that could not be explained by corresponding abundance changes alone.

**c**, PAE matrix for the AlphaFold 3-predicted mouse mitochondrial Complex V F1-containing assembly used to provide structural context for the BLIS PPI-remodeling results (WT; pTM = 0.78). Rows and columns correspond to aligned and scored residues, respectively, across the modeled assembly. Darker green indicates lower expected positional error and therefore greater confidence in the relative placement of the corresponding residues; lighter regions indicate greater positional uncertainty. Black lines delineate modeled chain or subunit boundaries. The extensive low-PAE blocks within and between major components support substantial confidence in the relative organization of large parts of the modeled assembly, while lighter off-diagonal regions identify interfaces or relative subunit orientations predicted with lower confidence. The model was used as structural context for the BLIS-remodeled ATP synthase interactions, which span the F1 α/β catalytic head, central-stalk and peripheral-stalk regions, rather than as a definitive experimental structure of Complex V.

Relate to **Fig.4f.**

**Extended Data Fig. 8 Conceptual comparison of the temporal weighting of molecular-state capture by direct fresh-tissue XL-MS and BLIS-XL-MS.**

**a,** Schematic representation of molecular-state sampling during direct crosslinking of freshly dissected tissue. Immediately after dissection ($t_{\mathrm{ini}}$), the tissue is depicted as being dominated by the initial native-like molecular-state ensemble (blue). During the interval required for crosslinker diffusion, reaction and quenching, continued *ex vivo* molecular remodeling can progressively alter protein conformations and interactions, increasing the contribution of remodeled states (magenta). At an illustrative intermediate time ($t_{x}$), both initial and remodeled states contribute to the population available for crosslinking. Because individual crosslinks can be formed throughout the interval between crosslinker addition and quenching ($t_{\mathrm{end}}$), the resulting XL-MS dataset represents a time-integrated molecular-state readout, rather than an instantaneous snapshot at the time of dissection. The purple bracket denotes the molecular-state accumulation window over which crosslinks formed at different times may contribute to the final measurement.

**b,** Conceptual representation of molecular-state sampling in BLIS-XL-MS. PFA fixation is used as the primary state-stabilization step, shifting the principal molecular-state capture window to the interval between the initial tissue state ($t_{\mathrm{ini}}$) and fixation-mediated state arrest ($t_{\mathrm{arrest}}$). Fixation is not instantaneous, and some *ex vivo* remodeling can therefore occur before molecular-state arrest. After fixation, cryosectioning and permeabilization shorten the diffusion path and improve reagent accessibility, enabling subsequent secondary crosslinking reactions to proceed in the stabilized tissue. Consequently, continued molecular remodeling during the longer secondary-crosslinking interval contributes substantially less to the final XL-MS readout than in direct fresh-tissue crosslinking. The solid purple bracket denotes the primary molecular-state accumulation window before state arrest, whereas the dashed interval indicates the reduced contribution of subsequent *ex vivo* remodeling during sample processing and secondary crosslinking. The final BLIS-XL-MS measurement therefore represents a fixation-stabilized, time-integrated structural and interaction fingerprint, rather than an unaltered atomic representation of the native conformational ensemble.

Blue and magenta curves denote the relative occupancy of initial native-like and *ex vivo*-remodeled molecular states, respectively. Protein cartoons illustrate representative states contributing to crosslink formation. Curves, crossover points and relative occupancies are schematic and are not intended to represent measured kinetic trajectories or quantitative state fractions.
